# Mapping Minds to Manage the ‘Tiger of Rivers’: A Fuzzy Cognitive Mapping Approach to Understanding Multi-Stakeholder Perspectives on Mahseer Conservation

**DOI:** 10.64898/2025.12.07.692881

**Authors:** Prantik Das, V. V. Binoy

## Abstract

Conservation of mahseers, a group of charismatic freshwater fishes native to the South, East and South-East Asia, situated at the intersection of complex social, ecological and economic landscapes, necessitates the integration of the knowledge, mental models, and expectation of different stakeholders. Using a Fuzzy Cognitive Mapping approach, this study generated collective mental models of mahseer conservation in two Indian states distinct in their geography, ecology and socio-cultural aspects - Assam and Uttarakhand. Although many core system components were similar in the cognitive map generated for the focal states, notable divergences emerged in the drivers, perceived relationships among the components, etc. indicating regional specificity in socio-cultural governance structures and stakeholder priorities. In both states ‘stakeholder communication’ emerged as the most central, and the component with highest outdegree, underscoring its shaping effect on system-wide variable interactions. The other influential components were ‘community fishing (destructive and traditional)’ and ‘human-human conflict’ (in Assam), and ‘illegal destructive fishing’ and ‘community involvement in decision-making’ (in Uttarakhand). The ‘what-if’ scenario analyses simulating potential interventions, demonstrated that enhancing communication, awareness and education, promoting local identity and cultural significance of community fishing, providing subsistence fishing opportunities for the locals, and community involvement in decision-making could help in reducing human-human conflicts, illegal fishing, and strengthen stakeholder collaboration for mahseer conservation in both states. Our results also indicate that these improvements alone are insufficient to result in measurable mahseer conservation outcomes until cultural revitalisation and governance reforms are integrated with habitat restoration, improved hatchery performance, evidence-based policies and strict law enforcement. We discuss various leverage points that emerged from the FCMs of both focal states and their implication for a broader mahseer conservation policy with a scope for integrating region-specific characteristics.

## Introduction

‘Tiger of rivers’ - as commonly known, mahseers are a group of iconic, freshwater fish species with many growing above 30 kg (known as megafishes). Inhabitants of the rivers of south, south east and east Asia, these fishes offer diverse religious, cultural, ecological, ecosystem and economical services. However, both natural populations and habitats of mahseers in many of these nations face a range of threats such as environmental degeneration, blockage of migration route due to dam construction, siltation, water pollution, destructive fishing methods (dynamiting, electrocution, poisoning, poaching, overfishing, destructive community and mass fishing), introduction of Invasive Alien Fishes (IAF), unregulated non-catch and release angling, etc. (Raghavan et al. 2011; Bhatt and Pandit 2016; Lewin et al. 2019). Although multiple schemes and policies have been implemented by many governmental organisations independently and in association with non-governmental actors, in their distribution range many mahseer species are continuing endangered or data deficient (Dahanukar et al. 2018; Pinder et al. 2019; Sarma et al. 2022). Furthermore, research focusing on this important group of freshwater fishes is dominated by biological, genetic, ecological and taxonomic dimensions (Malik 2011; Nautiyal 2001; Khare et al. 2014; Bhatt and Pandit 2016; Laskar et al. 2018; Pinder et al. 2019; Sarma et al. 2022; Dhawan et al. 2024) often overlooking the human aspects of conservation. However, it is well established that solving conservation challenges requires a deeper understanding of the mindset, values and actions of the stakeholders along with the biological information of the focal animals and the environmental data of their habitat (Collins et al. 2011). Unfortunately very few studies, mostly focused on a single stakeholder group recreational anglers (Gupta et al. 2014; 2015a; 2016a; 2016b; Bower et al. 2017), are available exploring the political, socio-cultural, and policy landscape within which the mahseer conservation occurs (Nautiyal 2014; Gupta et al. 2015b; Baruah 2024). However, knowledge of the perceptions, attitudes and patterns of behaviour (Banerjee et al. 2025; Prakash et al. 2025) by different stakeholders involved is vital to develop region-specific conservation strategies with ensured community involvement for successful implementation of these plans, which are generally long term in nature (Thant et al. 2023; Banerjee et al. 2025; Prakash et al. 2025).

Mapping the mindsets of the stakeholder individuals and groups, differing in their values, culture, reasoning style, associations, identities and norms and comprehensively converging them together for guiding locally inclusive evidence-based conservation policy and decision making (Nyboer et al. 2025) is a mammoth task. In this regard studying mental models, defined as personal and internal constructs which provide interpretation and structure of the environment and reality that people use to make sense and interact with the world around them (Craik 1943; Özesmi and Özesmi 2004; Jones et al. 2011; Gray et al. 2013; 2015; 2019) is getting lots of academic attention all over the world (Li et al. 2016; Cleveland et al. 2024; Segura et al. 2024; Banerjee et al. 2025; Prakash et al. 2025). Mental models are important components of an individual’s cognition, reasoning, decision making and also their personal assumptions about various socio-environmental issues (Moon et al. 2019). Individual mental models aggregate into a shared community mental model that functions as the foundation for collective cognition, collective intelligence and collaborative decision-making (Langan-Fox et al. 2001; van Velden et al. 2020; Blewett et al. 2021; 2022). Such a shared mental model also holds the potential to influence actions of both individuals and communities towards their external world (Özesmi and Özesmi 2004). Therefore, understanding and managing mental models existing at individual and community level is vital to find sustainable solutions for “wicked” environmental problems such as conservation of natural resources and the management of wildlife population requiring community-driven multi-actor, interventions (Gray et al. 2015; 2019; Li et al. 2016; Cleveland et al. 2024). However mental models are not static, and they shift with changing ecological, socio-cultural, and political contexts, and therefore require regular studies to track their ever-evolving attributes.

Fuzzy-logic Cognitive Mapping built on the concept of cognitive maps - mental representations of specific objects, events or concepts (Axelrod 1976) - is a powerful approach to decrypt and integrate mental models (Kosko 1986; Gray et al. 2013). Although, cognitive maps were developed initially as binary models (0 or 1; yes or no; Axelrod 1976), the awareness that cause-effect relationships existing between the variables of a social and environmental systems are inherently non-linear and non-deterministic, and could be better represented as fuzzy approximations of influences over binary relationships led Kosko (1986) to develop fuzzy-logic cognitive maps (Özesmi and Özesmi 2003; 2004). Fuzzy Cognitive Maps (FCMs) offer a semi-quantitative model of the conception and framing of a particular issue or problem in question by an individual and hence offers the possibility to combine expert knowledge and stakeholder perceptions into a holistic representation. FCM usually represents the knowledge by defining three characteristics viz., (i) the components/variables/factors present in a system, (ii) relationships (positive or negative), (iii) and the degree of influence existing between these components. To create the maps components of the focal system are described and the directed pairwise associations (positive or negative) are assigned. These associations (called edges) are allocated weight qualitatively (low, medium, high) or quantitatively (-1 to +1), to develop into a fuzzy dynamic model (Gray et al. 2013; 2015; 2019; Cleveland et al. 2024; Segura et al. 2024). FCM provides structural metric analysis (static) and dynamic, ‘what-if’ scenario analysis. The product from the former include graph theory indices such as the number of components/concepts (N), connections (C), transmitter (driver), receiver, ordinary, centrality, C/N, density and hierarchy index (Supplementary Materials SM 1; Özesmi and Özesmi 2003; Gray et al. 2014). Meanwhile the ‘what-if’ scenario analysis is useful in predicting the outcomes of the changes in any component (s) of the system (Özesmi and Özesmi 2004; Gray et al. 2019).

In the recent past FCM has evolved as a valuable tool in the field of conservation (Banerjee et al. 2025; Prakash et al. 2025). Facilitation of the insights on the points of consensus in the knowledge kept by different stakeholders and the mental models of the focal species and ecosystem they share, possibility to compare and combine these information (Bosma et al. 2017), provision of the potential trajectories that planning decisions and dynamics adaptational strategies (Bakhtavar et al. 2021) may take, makes fuzzy cognitive mapping a dependable tool for testing the feasibility of conservation interventions and policies (Knox et al. 2023). Furthermore, the potentials for integrating local knowledge into conservation management strategies and holistic visualisation of the complexity and interconnectedness between the between the actors the socio-ecosystems (Pluchinotta et al. 2019; Rooney et al. 2023) under focus enables FCM to perform the role of decision support mechanism (Papadopoulos et al. 2025) and augmenter of the communication efficacy in conservation (Hare 2011). Hence, in the recent past FCM received popularity in various fields aiming to find sustainable solutions for the wicked problems such as hydrology (Anezakis et al. 2016), environmental policy making (Kontogianni et al. 2012), management of bushmeat hunting and trade (Nyaki et al. 2014; van Velden et al. 2020), wildlife species and conflict management (Banerjee et al. 2015; Blewett et al. 2021; Prakash et al. 2025), agricultural policy (Vuillot et al. 2016), fisheries (Lavin et al. 2018) and water resource planning and management (Papadopoulos et al. 2025), etc.

Even after having immense potential, a sector which did not witness an FCM based approach yet is the mahseer conservation. Due to the complexity of social-ecological systems (SES) in which such activities are being implemented - freshwater ecosystems of 17 nations differing in their culture, including the trans-boundary rivers - support from FCM to visualise mental models of mahseers and their conservation and the relationship existing between numerous stakeholders, integrate their perspectives divergent in the values attributed to these migratory fishes, and identify key drivers, feedback loops, and leverage points would enhance the success of such interventions. A quick exploration of the literature will reveal the complexity associated with mahseer conservation; in Pakistan mahseer is the National Fish, they are religiously revered in selected regions of South-East Asia, while angled for recreation and eaten as a delicacy in some other areas of this continent (Philipp et al. 2015; Pinder et al. 2019; Muchlisin et al. 2022; Redhwan et al. 2022). The migratory route taken by many mahseers cut across the riverine landscapes divergent in the values and meanings attributed to them, and even cross political boundaries of nations which manifold the complexities of the situation. Hence, FCM could help in converging knowledge, emotions and expectations of important stakeholders, including national and regional governments, various departments, researchers, anglers, local communities, fish farmers, local NGOs, aquaculturists, and caretakers of temple/shrine sanctuaries to come up with sustainable mahseer conservation programmes and policies. Considering this scenario, the current study aimed to develop an FCM of mahseer conservation keeping two Indian states, Assam and Uttarakhand, known for their mahseer populations (Everard and Kataria 2011; Nautiyal 2014; Baruah 2017; Baruah and Sarma 2018; Pinder et al. 2019), in the focus. The current study aims to

1. Understand the components constituting collective mental models of mahseer conservation and the nature of relationship amongst them in the states of Assam and Uttarakhand?
2. Trace out leverage points with the potential to shape system-wide positive changes in mahseer conservation in the focal states?
3. Simulate intervention scenarios inspired from the result of the static analysis of FCMs to gain insights for improving mahseer conservation policy and management strategies

## Methodology

### Study Areas

The current study was conducted in two Indian states: Assam located in the Eastern, and Uttarakhand in the Western Himalayas. In Assam, mahseers are (mainly *T. putitora* and *N. hexagonolepis*) associated with recreational angling-based eco- and aqua-tourism activities that provides livelihoods and employment opportunities for the villagers (e.g. Nameri National Park area), age-old traditional community fishing festival such as *Junbeel Mela* (Baruah 2017; Baruah and Sarma 2018; Das et al. 2023), and have been traditionally fished for generations, being considered a rich source of protein and fetching high market prices (Baruah and Sarma 2018; Debnath et al. 2024). Here, breeding these species are supported by two hatcheries (Borgohain 2015; Das and Gogoi 2015), but illegal fishing, poaching, destructive community fishing practices, habitat degradation and breeding ground destruction through dam construction, siltation have intensified pressures on the wild mahseer populations inhabiting rivers and *beel*/wetlands or lakes (Medhi and Sharma 2017; Dutta et al. 2023). Frictions between local community members and government officials related to various community fishing-related activities have also increased in the recent past (Mitra 2015; Bhattacharjee 2025) We collected data from four districts viz., Kamrup (Metropolitan), Morigaon, Nagaon and Sonitpur (Fig. 1) from this state.

**Fig 1.**
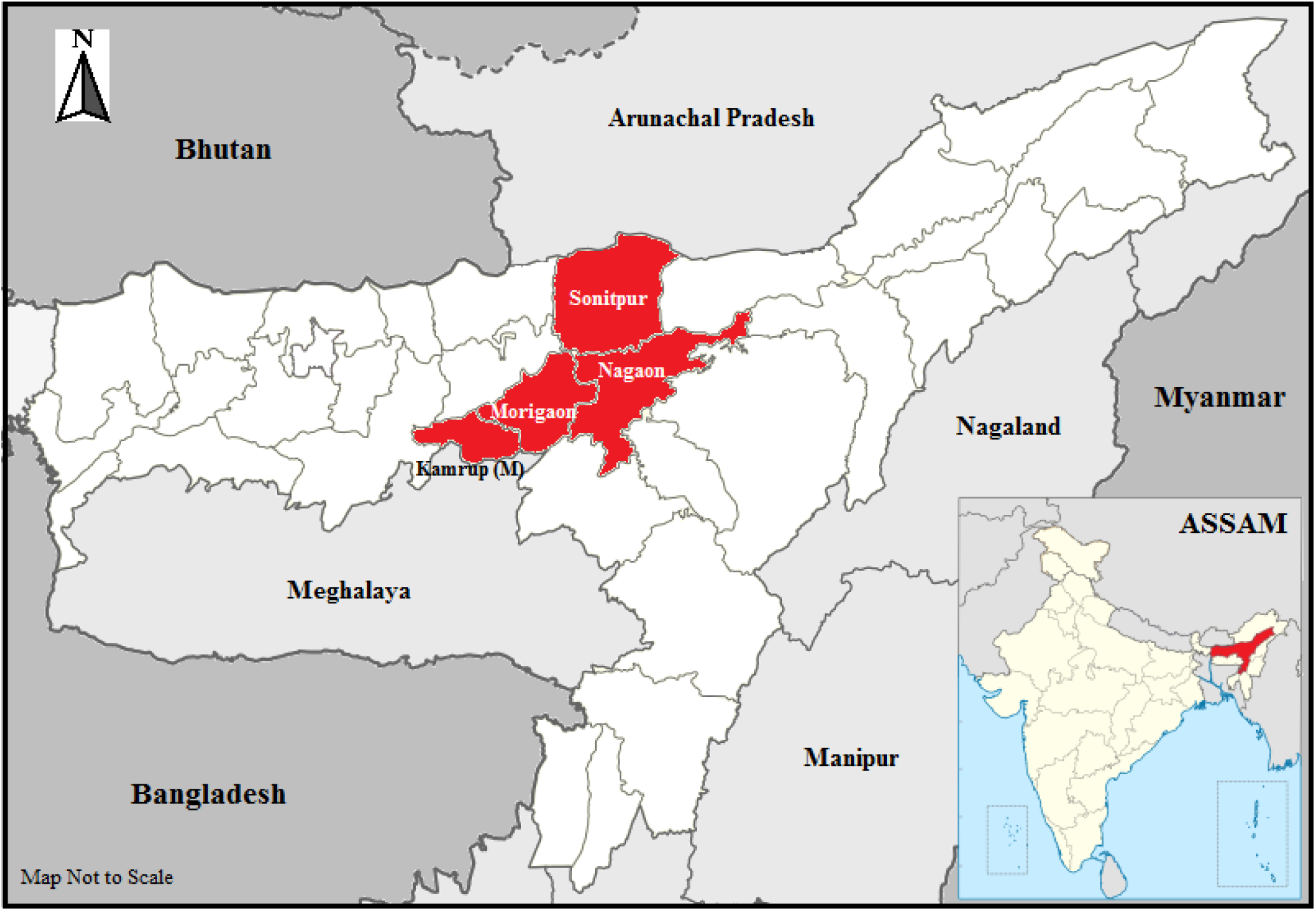
The districts of the Indian state of Assam where the field study was conducted. Map not to scale.

In Uttarakhand, mahseer (*T. putitora*) holds the special position of State Fish, with wild populations replenished by ICAR–DCFR hatchery (ICAR 2016). This state is an internationally renowned destination for the recreational angling of mahseer, particularly the Pancheshwar region, where eco-tourism provides significant local livelihood opportunities. Traditional community fishing practices and rituals such as the *Maund* (or *Maun*) *Matsya Mela,* which are undergoing commercialisation and witnessing excessive usage of unsustainable fishing methods leading to the decline of mahseer population are also reported from Uttarakhand (Sharma et al. 2016; Sundriyal and Kumar 2019; Uniyal and Uniyal 2021). At the same time many regions of this state have temple sanctuaries such as Girjiya Devi and Baijnath temple, where these fishes are revered and protected. Alongside these, a variety of other activities involving multiple stakeholders such as recreational angling and eco-tourism based benefit sharing, hydropower and highway development projects, river-mining and destructive community fishing-related events, have often been shown to generate conflicts arising from divergent competing stakeholder interests (Bower et al. 2017; Jain 2020; Tapasya 2024; Azad and Talwar 2025; Sharma 2025). Dehradun, Tehri Garhwal, Nainital and Champawat (Fig. 2) were the four districts chosen for the fieldwork.

**Fig 2.**
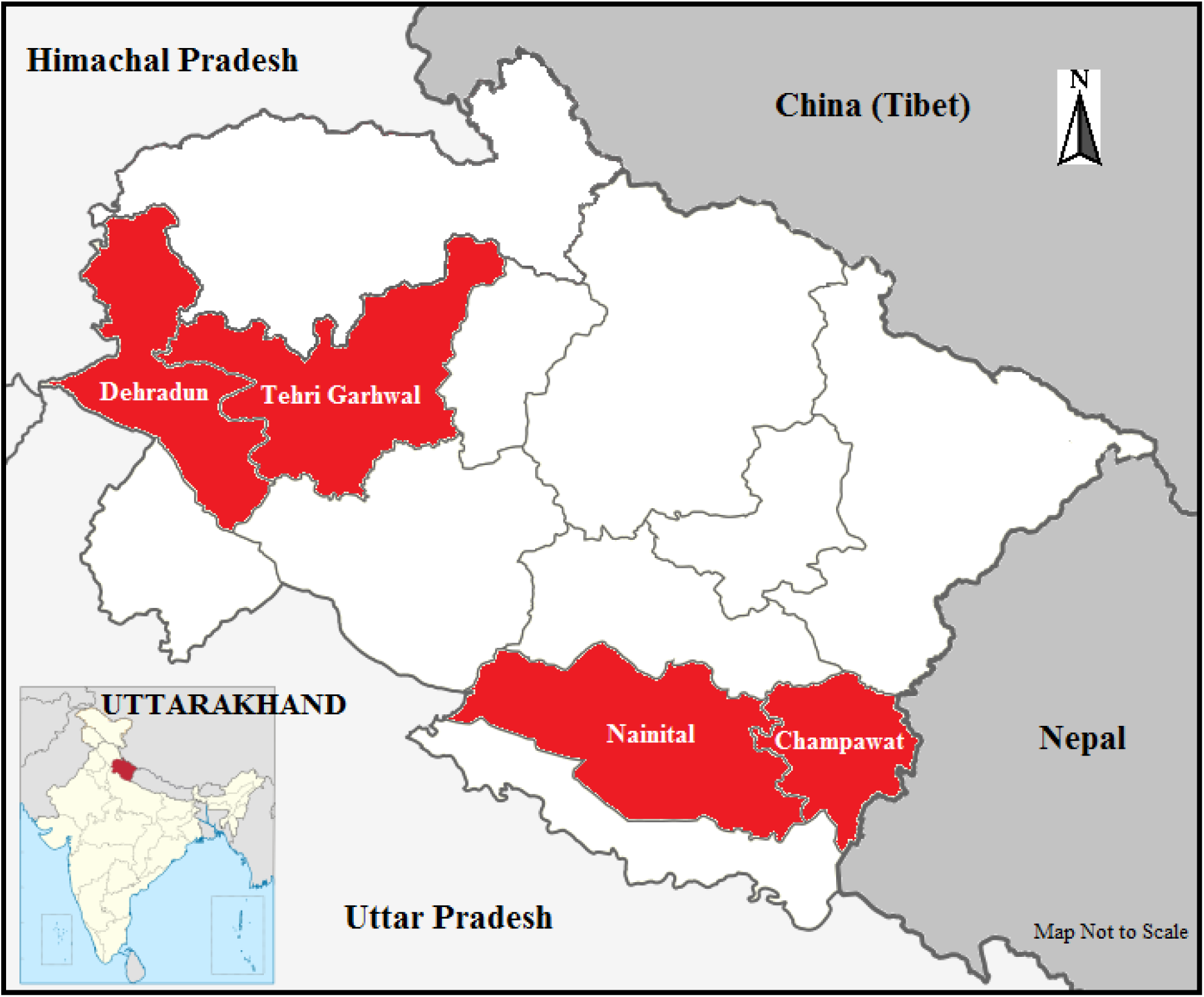
The districts of the Indian state of Uttarakhand where the field study was conducted. Map not to scale.

### Data Collection and Analysis

From each focal state, we tried to collect information from a diverse group of stakeholders (Table 1) possible. Snowball sampling was used for identifying the key individuals and groups who could provide rich information on mahseer. The individual mapping, which provides a robust and nuanced representation of the participants’ understanding and reveals individual differences existing in the concepts, knowledge and perception (Gray et al. 2014) was the technique used to generate the data required for fuzzy cognitive mapping. This approach is advantageous over facilitated group modelling, another methodology popular in FCM, due to the equitable and individual-level representations of the knowledge it provides (Özesmi and Özesmi 2004; Gray et al. 2014; Banerjee et al. 2025). In-depth, discursive, semi-structured one-on-one interviews (for questionnaire - Appendix 1) of the interested participants, duration of which ranged between 15 to 45 minutes, were conducted to trace out their mental models of mahseer conservation and behaviours. Some stakeholder groups expressed a preference for group discussions instead of individual interviews. While conducting the focus group discussions (FGDs), to reduce the influence of any external factors, the questions were asked to each individual participant within the group-setting. We conducted 91 interviews and 2 FGDs (number of participants = 11 and 12) in Assamese, English and Hindi languages in Assam and 44 interviews (Hindi) in Uttarakhand (Table 1).

**Table 1.**
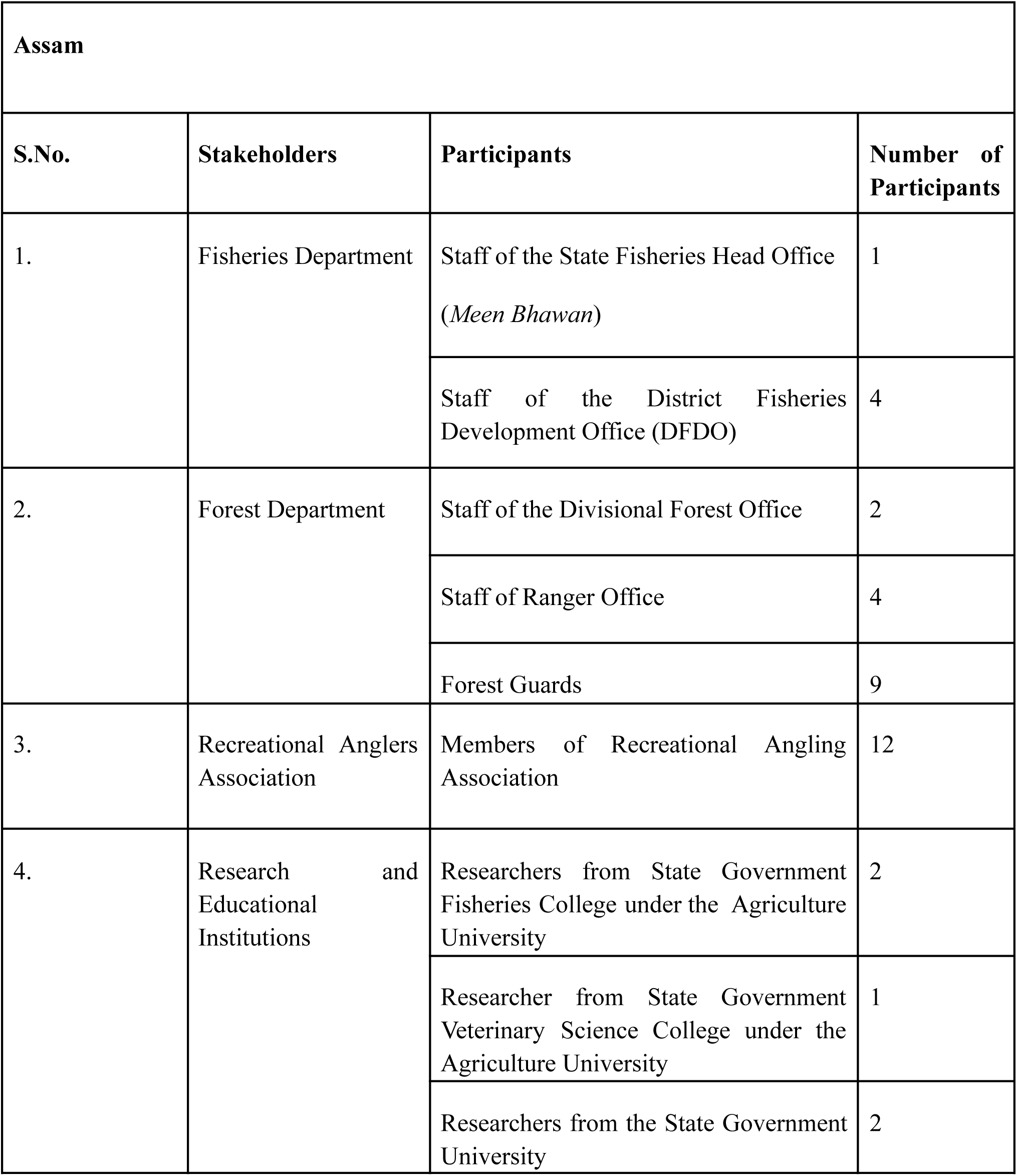
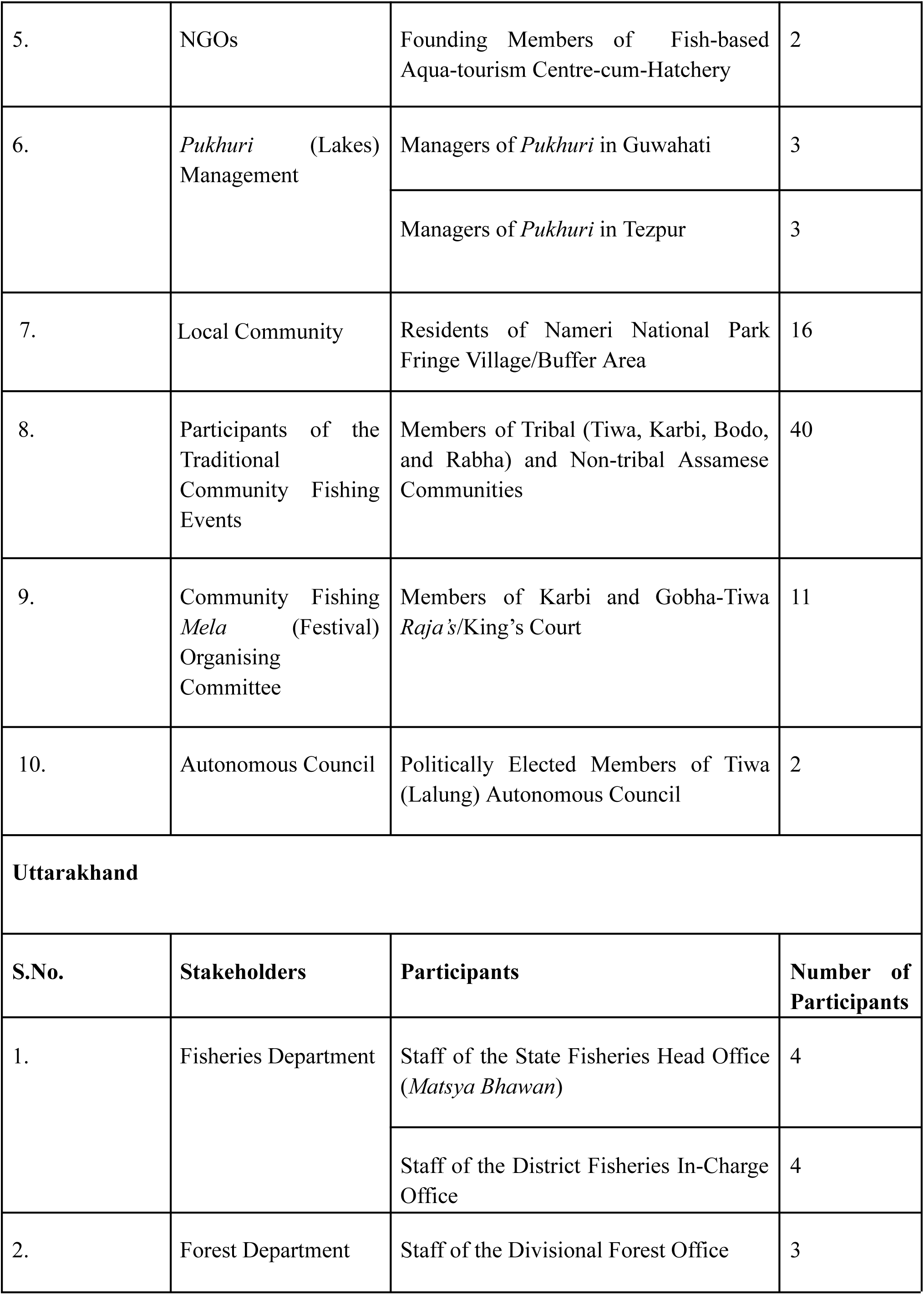
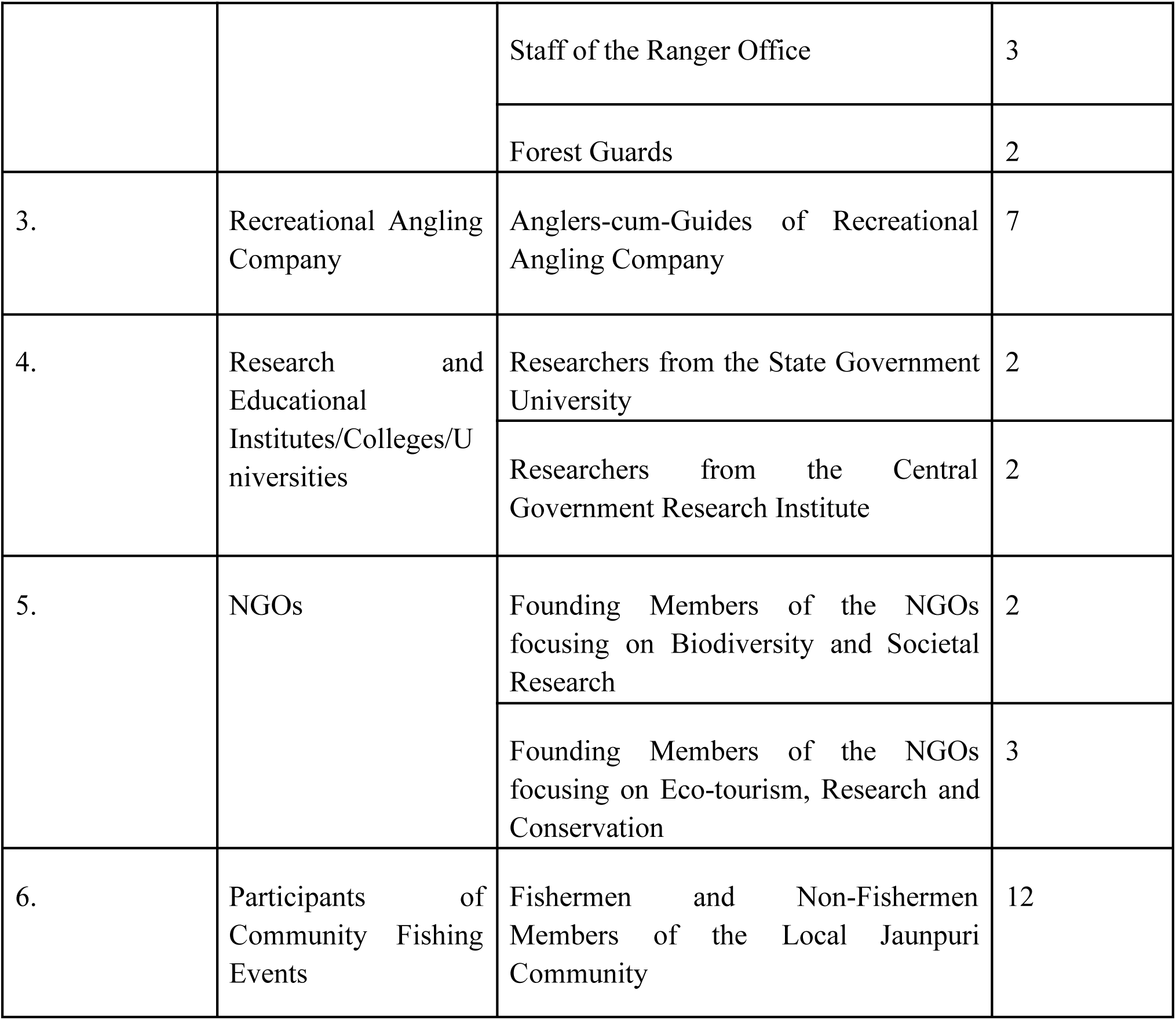
List of stakeholders from Assam and Uttarakhand interviewed/participated in the Focus Group Discussion (FDG).

To establish components/variables (parameters linked to the mahseer conservation) present in their mental models (Salberg et al. 2022) for FCM the indirect elicitation with free association of concepts method (Gray et al. 2012; 2014) was followed. Here participants were allowed to freely choose their own concepts regarding the mahseer and their conservation in response to the question from the researcher. The variables and the strength and direction of the relationships between them, emerged from the answers of the participants, were used for generating individual FCMs. For this purpose interviews in English were transcribed verbatim using Otter.ai and Rev.ai. The responses in vernacular languages were manually transcribed and translated into English. All transcripts were carefully cross-checked against the original audio recordings and field notes were reviewed to avoid ambiguities. The final version of the transcripts were then used for the extrapolation and construction of individual FCMs by the first author (PD) using the ‘Mental Modeler’ (MM; http://www.mentalmodeler.org; Gray et al. 2013; 2017). Tools such as MM develop qualitative FCMs which are then translated into semi-quantitative structures required to run dynamic future scenarios (Gray et al. 2013) by artificially increasing or decreasing the values of the system components (known as clamping; Özesmi and Özesmi 2004; Gray et al. 2013; 2015; 2019). We chose MM for the current study due to its user-friendly interface, participatory focus, specific design for multi-stakeholder engagement and model building, real-time, dynamic graphical outputs that improve comprehension, and built-in analytical functions such as centrality, outdegree, and indegree to identify leverage points and key driving components of the system, etc. (Gray et al. 2013, 2015). Additionally MM has been effectively used in conservation, nature resource management and policy planning by various authors (Nyaki et al. 2014; Li et al. 2016; Gray et al. 2019; van Velden et al. 2020; Blewett et al. 2021; Segura et al. 2024).

Edge relationships between the components were then mapped into individual adjacency matrices. The relationships between different components that were qualitatively defined by the respondents (low, medium and high) were subjected to fuzzy scoring. The ‘low, medium and high’ attributed by the respondents were converted to 0.25, 0.50 and 1 respectively, with negative relationships encoded as inverse, and 0 representing no relationship (Özesmi and Özesmi 2003; 2004; Nyaki et al. 2014; Henley-Shepherd et al. 2015; Li et al. 2016; van Velden et al. 2020). A total of 114 adjacency matrices representing individual cognitive maps for the state of Assam and 44 for the Uttarakhand was generated following this methodology. The individual adjacency matrices were superimposed to create a shared or collective cognitive map for each of the two focal states by taking the average of the weighted edge relationships (SM 2a, 2b; Özesmi and Özesmi; Gray et al. 2012; Li et al. 2016). Here equal weightage was attributed to individual maps to ensure their validity and importance (Li et al. 2016; van Velden et al. 2020), as well as to prevent stakeholder groups with a higher number of representatives from exerting a disproportionate influence on the overall community map (Salberg et al. 2022). The collective FCMs generated for Assam and Uttarakhand were subjected to structure (structural and static analysis of the graph theory indices), content (the extent of shared or unique concepts) and “what-if” scenario analysis for the comparison (Yoon and Jetter 2016; Blewett et al. 2021).

The total number of components, connections, density, connections per component, number of driver, receiver and ordinary components, hierarchy score, and the indegree, outdegree and the centrality value of each variable in the map (Table 2, 3; Özesmi and Özesmi 2004) were compared. Map similarity indices viz., S2 (the proportion of shared concepts between maps; 0 = completely dissimilar; 1 = identical structure; SM 1) and Jaccard (for weighted adjacency matrix of FCM; a ratio of shared versus unique edge relationships between components; 0 = no shared edges; 1 = identical edges; SM 1) indices were estimated (van Velden et al. 2020).

**Table 2.** Structural metrics of collective Fuzzy Cognitive Maps (FCMs) of Assam and Uttarakhand.

| **Structural Metric** | **Assam FCM** | **Uttarakhand FCM** |
| --- | --- | --- |
| Total Components | 44 | 42 |
| Total Connections | 166 | 185 |
| Connections per component | 3.772 | 4.404 |
| Density | 0.09 | 0.11 |
| Number of driver components | 9 | 1 |
| Number of receiver components | 0 | 0 |
| Number of ordinary components | 35 | 41 |
| Hierarchy Index | 0.000128 | 0.000301 |
| Clustering Coefficient (Global) | 0.28 | 0.36 |

For the “what if” analysis, we identified key components, which can act as ‘leverage points’ capable of inducing system-wide ripple effects, to construct alternative scenarios (Meadows 1997; Meadows and Wright 2008; Gray et al. 2015; Banerjee et al. 2025; SM 1). Major drivers, components with the high centrality and outdegree values, and participant recommendation factors (Salberg et al. 2022), were used for determining these leverage points (Kosko 1986; Özesmi and Özesmi 2004; Gray et al. 2015; 2019; Li et al. 2016; van Velden et al. 2020; Blewett et al. 2021; 2022; Salberg et al. 2022; Segura et al. 2024). Responses to the question, *“What additional recommendations would you like to suggest to shape a potential future system that benefits both the mahseers and associated stakeholders?,”* functioned as the basis for the latter. We then ran what-if simulations by clamping the selected components as well as the recommendations, using a sigmoid activation and stabilising function to analyse the resulting effects on the steady state of the FCM (Bueno and Salmeron 2009). The outcome of these scenarios were subsequently compared to the original steady state to assess the changes occurring in the components in response to the modifications of the selected components (Gray et al. 2015). We used Mental Modeler for static, structural and scenario analyses, FCMapper package function comp.maps for the content comparisons, Igraph and matrix.indices to compute clustering coefficient (global) and hierarchy indices for the structure analyses, and VennDiagram and ggplot2 packages for visualisations in RStudio versions 2024.09.0 and 2025.05.0.

## Results

### Structural/Static Analysis

The shared collective FCM of Assam comprised 44 components interconnected by 166 connections (Fig 3), while the Uttarakhand FCM had 42 components with 185 connections (Fig 4). However, Assam’s map had a higher number of drive variables (9), as compared to that of Uttarakhand (drive variable = 1), while the map densities of both these FCMs were not much different (Uttarakhand; U: 0.11; Assam; A: 0.09; Table 2). “Socio-cultural significance”, “social media for awareness”, “community involvement in decision making”, “subsistence fishing for local community members”, “species breeding and biological barriers”, “limited time availability”, “promotion of Assamese and Tribal culture and identity”, “promotion of folklores and songs related to community fishing” and “development of *beel* for community fishing” were drivers emerged for Assam. Interestingly, only one driver variable (“declaration of State Fish”) was found in the Uttarakhand FCM. However, because of the low outdegree values (0.3; Table 3) this component (declaration of State Fish) was considered only as a micro-variables and not a major driver (Banerjee et al. 2025). Such micro-variables observed in Assam FCM with low outdegree values were: “social media for awareness (0.04)”, “species’ breeding and biological barriers (0.04)”, “limited time availability (0.02)” and “development of *beel* for community fishing (0.92)”. The clustering coefficient (global) obtained for Uttarakhand (0.36) was slightly greater than Assam (0.28), and both states received a hierarchy index near to zero (A: 0.000128; U: 0.000301; Table 2). Additionally, no receiver variables were found to be present in both states. Although ‘mahseer conservation’ was the most central variable for both states (centrality - A: 13.89, U: 19.92, Table 3, 4; indegree A: 12.51; U: 16.64) it was not considered for the comparison due to its status of ‘system focus’ (the dependent variable; Li et al. 2016; van Velden et al. 2020; Segura et al. 2024; Prakash et al. 2025).

**Fig 3.**
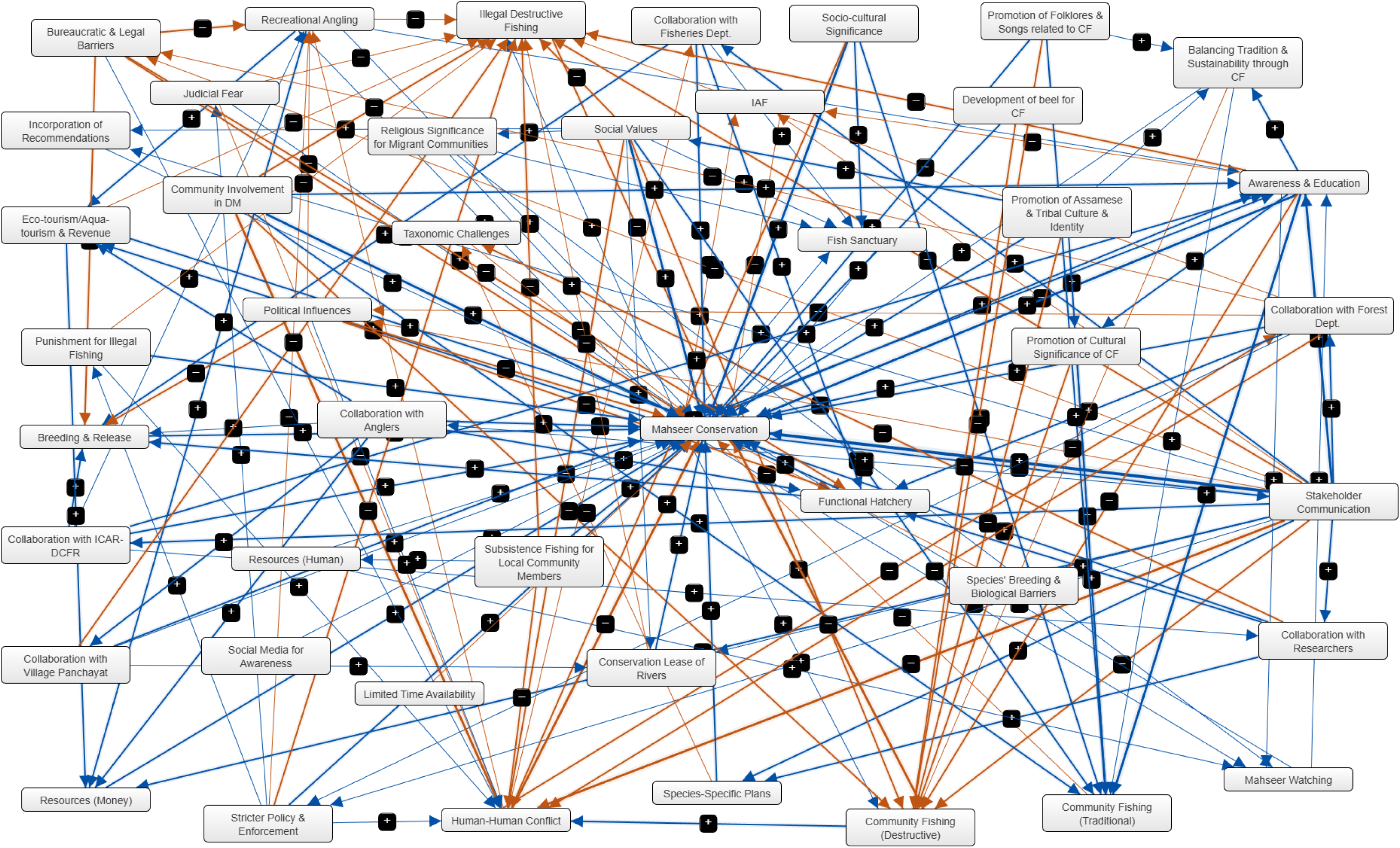
The collective Fuzzy Cognitive Map (FCM) generated by averaging the adjacency matrices of 114 respondents from Assam. CF: Community Fishing; DM: Decision-Making; IAF: Invasive Alien Fishes.

**Fig 4.**
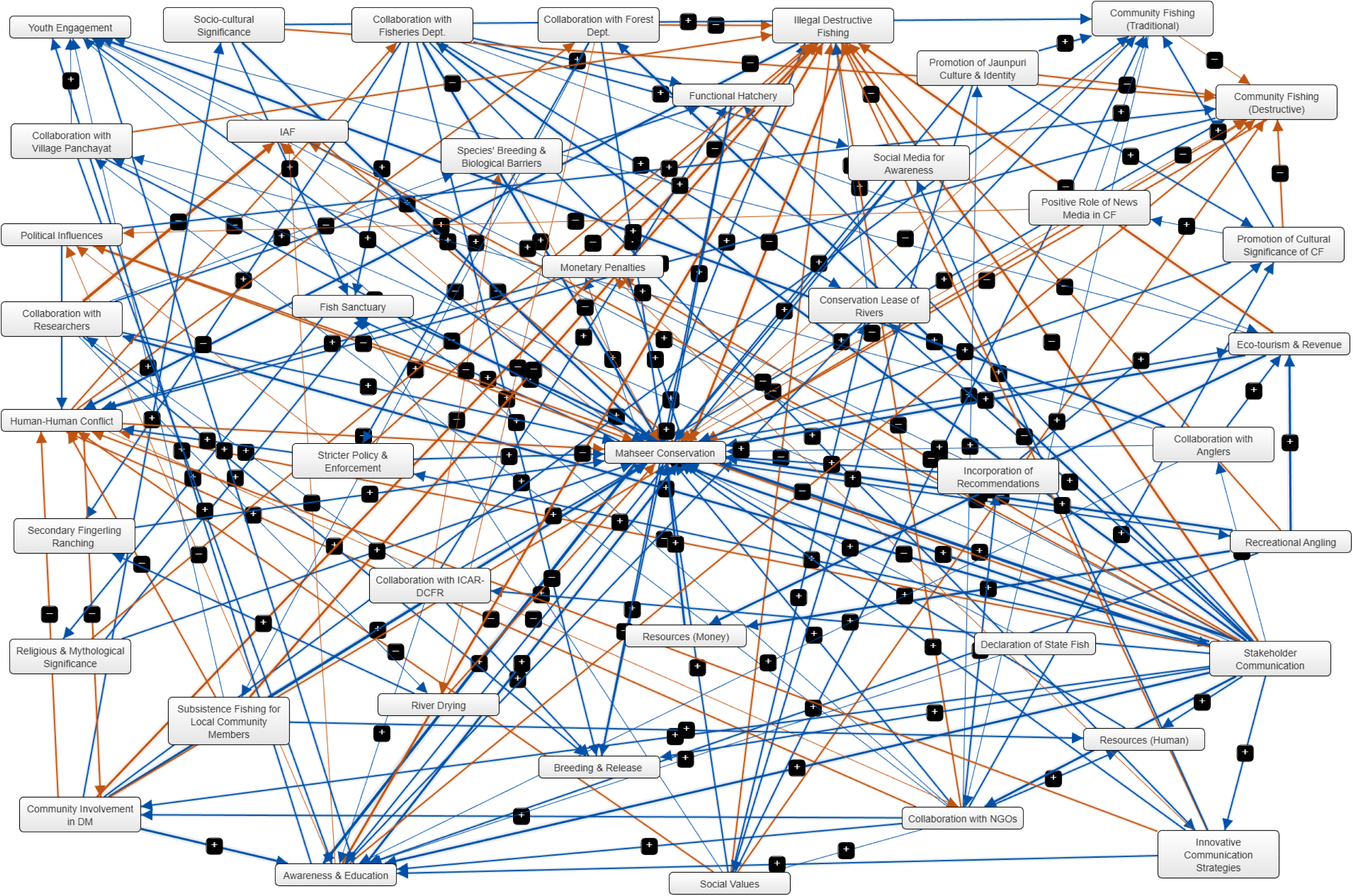
The collective Fuzzy Cognitive Map (FCM) generated by averaging the adjacency matrices of 44 respondents from Uttarakhand. CF: Community Fishing; DM: Decision-Making; IAF: Invasive Alien Fishes.

**Table 3.** Structural metrics of all components of the collective FCM of Assam.

| **Components** | **Indegree** | **Outdegree** | **Centrality** | **Component Type** |
| --- | --- | --- | --- | --- |
| Mahseer Conservation | 12.51 | 1.39 | 13.90 | Ordinary |
| Recreational Angling | 0.70 | 0.99 | 1.69 | Ordinary |
| Illegal Destructive Fishing | 2.49 | 0.32 | 2.82 | Ordinary |
| Community Fishing (Traditional) | 3.41 | 0.05 | 3.46 | Ordinary |
| IAF | 0.18 | 0.13 | 0.31 | Ordinary |
| Stakeholder Communication | 0.32 | 4.95 | 5.27 | Ordinary |
| Collaboration with Fisheries Department | 0.32 | 0.85 | 1.17 | Ordinary |
| Collaboration with Researchers | 0.36 | 0.72 | 1.08 | Ordinary |
| Breeding and Release | 1.34 | 0.31 | 1.65 | Ordinary |
| Resources (Financial) | 1.20 | 0.22 | 1.42 | Ordinary |
| Resources (Human) | 0.13 | 0.20 | 0.33 | Ordinary |
| Human-Human Conflict | 3.45 | 1.09 | 4.54 | Ordinary |
| Social Values | 0.46 | 2.36 | 2.82 | Ordinary |
| Socio-cultural Significance | 0 | 1.83 | 1.83 | Driver |
| Collaboration with ICAR-DCFR | 0.21 | 0.69 | 0.90 | Ordinary |
| Collaboration with Forest Department | 0.58 | 1.30 | 1.88 | Ordinary |
| Awareness and Education | 2.27 | 2.96 | 5.23 | Ordinary |
| Social Media for Awareness | 0 | 0.04 | 0.04 | Driver |
| Stricter Policy and Enforcement | 0.17 | 0.83 | 1.00 | Ordinary |
| Community Involvement in Decision Making | 0 | 3.01 | 3.01 | Driver |
| Subsistence Fishing Right for Local Community Members | 0 | 1.03 | 1.03 | Driver |
| Eco-tourism/Aqua-tourism and Revenue | 0.93 | 0.93 | 1.86 | Ordinary |
| Judicial Fear | 0.13 | 0.29 | 0.42 | Ordinary |
| Political Influences | 0.18 | 1.05 | 1.23 | Ordinary |
| Bureaucratic and Legal Barriers | 0.22 | 1.15 | 1.37 | Ordinary |
| Taxonomic Challenges | 0.21 | 0.15 | 0.36 | Ordinary |
| Species' Breeding and Biological Barriers | 0 | 0.04 | 0.04 | Driver |
| Fish Sanctuary | 0.69 | 0.10 | 0.79 | Ordinary |
| Conservation Lease of Rivers | 0.30 | 0.27 | 0.57 | Ordinary |
| Community Fishing (Destructive) | 3.91 | 1.09 | 5.00 | Ordinary |
| Functional Hatchery | 1.32 | 0.49 | 1.81 | Ordinary |
| Limited Time Availability | 0 | 0.02 | 0.02 | Driver |
| Promotion of Assamese and Tribal Culture and Identity | 0 | 2.08 | 2.08 | Driver |
| Punishment for Illegal Fishing | 0.13 | 0.55 | 0.68 | Ordinary |
| Incorporation of Recommendations | 0.04 | 0.02 | 0.06 | Ordinary |
| Collaboration with Anglers | 0.13 | 1.39 | 1.52 | Ordinary |
| Promotion of Cultural Significance of Community Fishing | 0.57 | 1.40 | 1.97 | Ordinary |
| Promotion of Folklores and Songs related to Community Fishing | 0 | 1.49 | 1.49 | Driver |
| Development of *beel* for Community Fishing | 0 | 0.92 | 0.92 | Driver |
| Balancing Tradition and Sustainability through Community Fishing | 0.68 | 0.33 | 1.01 | Ordinary |
| Mahseer Watching | 0.21 | 0.08 | 0.29 | Ordinary |
| Religious Significance for North-West Indian Migrant-origin Communities in Assam | 0.05 | 0.21 | 0.26 | Ordinary |
| Collaboration with Village Panchayat | 0.25 | 0.87 | 1.12 | Ordinary |
| Species-Specific Plans | 0.34 | 0.21 | 0.55 | Ordinary |

The ‘top 5’, and hence most influential central components (Gray et al. 2015) for Assam FCM were “stakeholder communication” (centrality value = 5.27), “awareness and education” (5.23), “community fishing (destructive)” (5)), “human-human conflict” (4.54), and “community fishing (traditional)” (3.46)) (Table 3). Among these “Community fishing (traditional)” (3.41)), “human-human conflict” (3.45), “community fishing (destructive)” (3.91) had the highest levels of indegree (≥ 3; van Velden et al. 2020), and “stakeholder communication” (4.95) and “community involvement in decision making” (3.01) the outdegree (≥ 3; van Velden et al. 2020). In Uttarakhand FCM, “stakeholder communication” (9.10), “illegal destructive fishing” (7.90), “awareness and education” (6.92), “human-human conflict” (4.74) and “community involvement in decision making” (4.25; Table 4) occupied the top 5 position. Here, “community fishing (destructive)” (3.08), “human-human conflict” (3.61), “illegal destructive fishing” (6.46) and “awareness and education” (4.30) had high levels of indegree. Meanwhile, the high levels of outdegree was associated with “stakeholder communication” (8.53) and “community involvement in decision making” (3.27). In both states the only central variable with a greater outdegree-to-indegree value was the “stakeholder communication” (A, 15.46; U, 14.96).

**Table 4.** Structural metrics of all the components of the collective FCM of Uttarakhand.

| **Components** | **Indegree** | **Outdegree** | **Centrality** | **Component Type** |
| --- | --- | --- | --- | --- |
| Mahseer Conservation | 16.64 | 3.28 | 19.92 | Ordinary |
| IAF | 1.37 | 0.30 | 1.67 | Ordinary |
| Social Values | 0.31 | 2.04 | 2.35 | Ordinary |
| Community Fishing<br>(Destructive) | 3.08 | 0.68 | 3.76 | Ordinary |
| Recreational Angling | 0.34 | 2.41 | 2.72 | Ordinary |
| Collaboration with<br>Fisheries Department | 0.75 | 2.21 | 2.96 | Ordinary |
| Collaboration with<br>NGOs | 0.79 | 2.50 | 3.29 | Ordinary |
| Collaboration with<br>Researchers | 0.67 | 2.13 | 2.80 | Ordinary |
| Stakeholder<br>Communication | 0.57 | 8.53 | 9.10 | Ordinary |
| Incorporation of<br>Recommendations | 0.20 | 0.77 | 0.97 | Ordinary |
| Community Involvement<br>in Decision Making | 0.98 | 3.27 | 4.25 | Ordinary |
| Breeding and Release | 1.52 | 0.43 | 1.95 | Ordinary |
| Resources (Financial) | 1.07 | 0.57 | 1.64 | Ordinary |
| Resources (Human) | 0.60 | 0.48 | 1.08 | Ordinary |
| Monetary Penalties | 0.37 | 0.95 | 1.32 | Ordinary |
| Human-Human Conflict | 3.61 | 1.13 | 4.74 | Ordinary |
| Fish Sanctuary | 1.43 | 0.99 | 2.42 | Ordinary |
| Subsistence Fishing for Local Community Members | 0.05 | 1.71 | 1.76 | Ordinary |
| Illegal Destructive Fishing | 6.46 | 1.44 | 7.90 | Ordinary |
| Collaboration with Forest Department | 0.75 | 0.87 | 1.62 | Ordinary |
| Collaboration with ICAR-DCFR | 0.27 | 1.20 | 1.47 | Ordinary |
| Eco-tourism and Revenue | 1.31 | 1.83 | 3.14 | Ordinary |
| Socio-cultural Significance | 0.39 | 1.93 | 2.35 | Ordinary |
| Religious and Mythological Significance | 0.27 | 0.83 | 1.10 | Ordinary |
| Collaboration with Anglers | 0.16 | 1.31 | 1.47 | Ordinary |
| Stricter Policy and Enforcement | 0.54 | 1.27 | 1.81 | Ordinary |
| Community Fishing (Traditional) | 2.38 | 0.29 | 2.67 | Ordinary |
| Awareness and Education | 4.29 | 2.63 | 6.92 | Ordinary |
| Promotion of Jaunpuri Culture and Identity | 0.11 | 1.52 | 1.63 | Ordinary |
| Social Media for Awareness | 0.37 | 0.94 | 1.31 | Ordinary |
| Political Influences | 0.43 | 0.60 | 1.03 | Ordinary |
| Species' Breeding and Biological Barriers | 0.20 | 0.16 | 0.36 | Ordinary |
| Conservation Lease of Rivers | 0.79 | 0.26 | 1.05 | Ordinary |
| Functional Hatchery | 0.99 | 0.70 | 1.69 | Ordinary |
| Promotion of Cultural Significance of Community Fishing | 0.97 | 1.32 | 2.29 | Ordinary |
| Collaboration with Village Panchayat | 0.38 | 1.10 | 1.48 | Ordinary |
| Positive Role of News Media in Community Fishing | 0.11 | 0.26 | 0.37 | Ordinary |
| Youth Engagement | 1.48 | 0.78 | 2.26 | Ordinary |
| River Drying | 0.15 | 0.66 | 0.81 | Ordinary |
| Secondary Fingerling Ranching | 0.37 | 0.18 | 0.55 | Ordinary |
| Innovative Communication Strategies | 0.37 | 1.10 | 1.47 | Ordinary |
| Declaration of State Fish | 0 | 0.30 | 0.30 | Driver |

### Content Analysis

We found 32 shared components in the FCMs of Assam and Uttarakhand, with a significantly high (S2 = 0.973; Fig 5) similarity in the components/variables. However, the links shared between these variables by the FCMs of two focal states were relatively low (Jaccard = 0.293; Fig 5). “Aqua-tourism and revenue”, “judicial fear”, “bureaucratic and legal barriers”, “taxonomic challenges”, “limited time availability”, “punishment for illegal fishing”, “promotion of folklores and songs related to community fishing”, “development of *beel* for community fishing”, “balancing tradition and sustainability through community fishing”, “mahseer watching”, “religious significance for North-West Indian migrant-origin communities in Assam”, “species-specific plans” were the variables unique to Assam (Fig 6). Distinctive variables that emerged in the Uttarakhand FCM were “collaboration with NGOs”, “monetary penalties”, “eco-tourism and revenue”, “religious and mythological significance”, “positive role of news media in community fishing”, “youth engagement”, “river drying”, “secondary fingerling ranching”, “innovative communication strategies” and “declaration of State Fish” (Fig 7).

**Fig 5.**
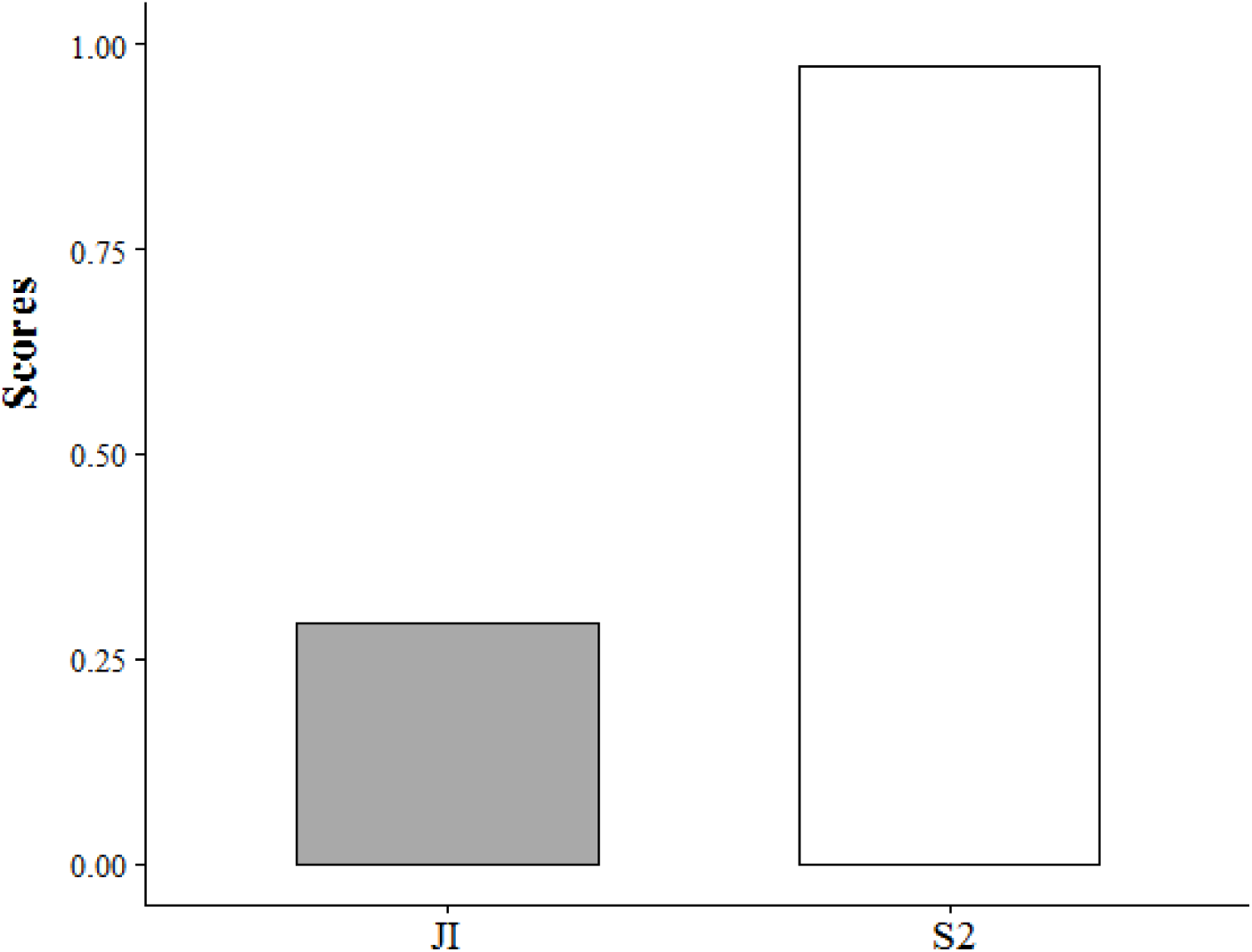
Jaccard (JI) and Similarity (S2) Indices indicating the similarity between the collective FCMs of Assam and Uttarakhand.

**Fig 6.**
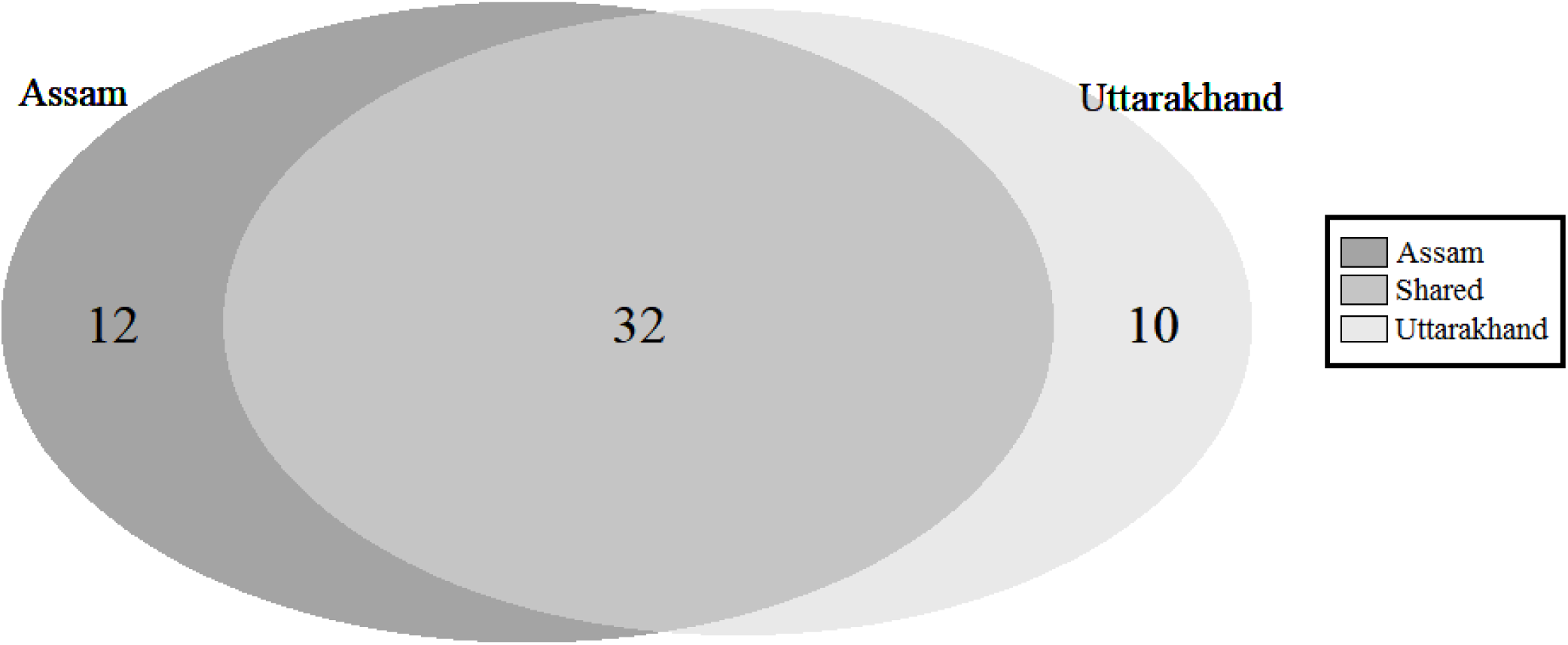
Unique and shared components in the collective FCMs of Assam and Uttarakhand.

**Fig 7.**
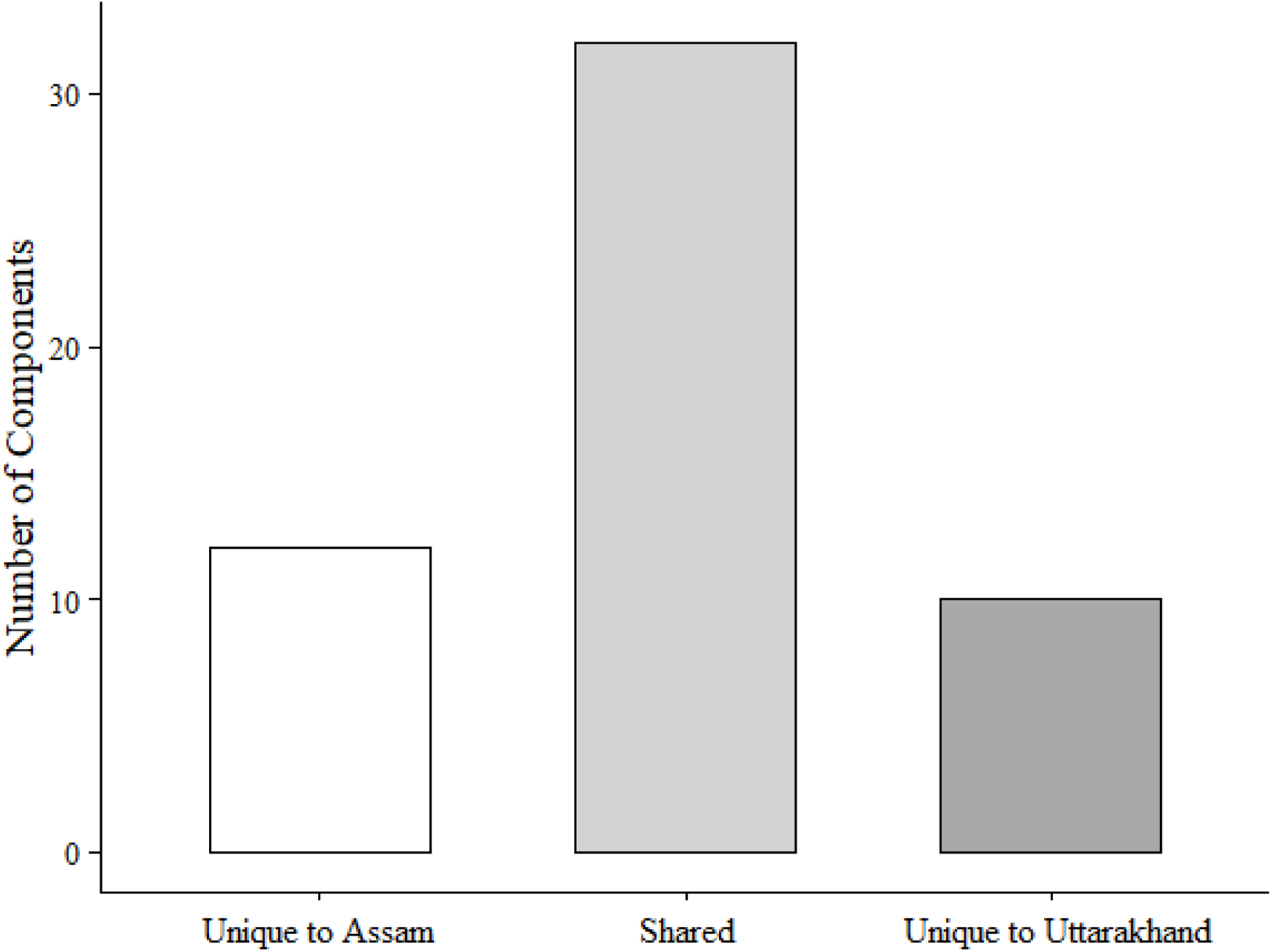
Unique and shared components in the collective FCMs of Assam and Uttarakhand.

### What-if Scenario Analysis

We tested four “what-if” scenarios representing different combinations of potential mahseer conservation interventions by simultaneously clamping multiple system components (drivers, components with high centrality and out-degree values) and stakeholder recommendations for both focal states. The criteria used for clamping was as follows; if a system component was further recommended by the stakeholders we clamped it +1 or -1. The negative or positive sign was attributed based on the increase or decrease as proposed by the stakeholders. However, clamping was restricted to +0.5 or −0.5 if the component was only a ‘stakeholder recommendation’ and did not fall under the above mentioned category of system components (Gray et al. 2019; van Velden et al. 2020; Salberg et al. 2022; Segura et al. 2024).

In the first scenario (Scenario 1; Fig 8a) tested in Assam, the following variables were increased to +1: stakeholder communication, awareness and education, community fishing (traditional), community involvement in decision making, subsistence fishing for locals, promotion of Assamese and Tribal culture and identity, promotion of folklores and songs related to community fishing and socio-cultural significance of mahseers. Meanwhile resources (money and human), and fish sanctuaries, were increased to + 0.5 only. Under this scenario, 60.60% (20 out of the 33) of the FCM components exhibited a trajectory shift while others remained in the steady state. Components such as illegal destructive fishing (−0.07), human-human conflict (−0.14) and community fishing (destructive; −0.07) declined, while a marginal decrease was observed in the IAF, political influences, bureaucratic and legal barriers (all −0.01; Fig 8a). The upward shift was seen in social values (0.06), balancing traditions and sustainability through community fishing (0.05), collaboration with forest department (0.04), promotion of cultural significance of community fishing (0.04), collaboration with fisheries department and researchers (both 0.03), collaboration with DCFR-ICAR and species-specific plans (both 0.02), breeding and release, stricter policy and enforcement, conservation lease of rivers, functional hatchery, collaboration with anglers (all 0.01).

**Fig 8.**
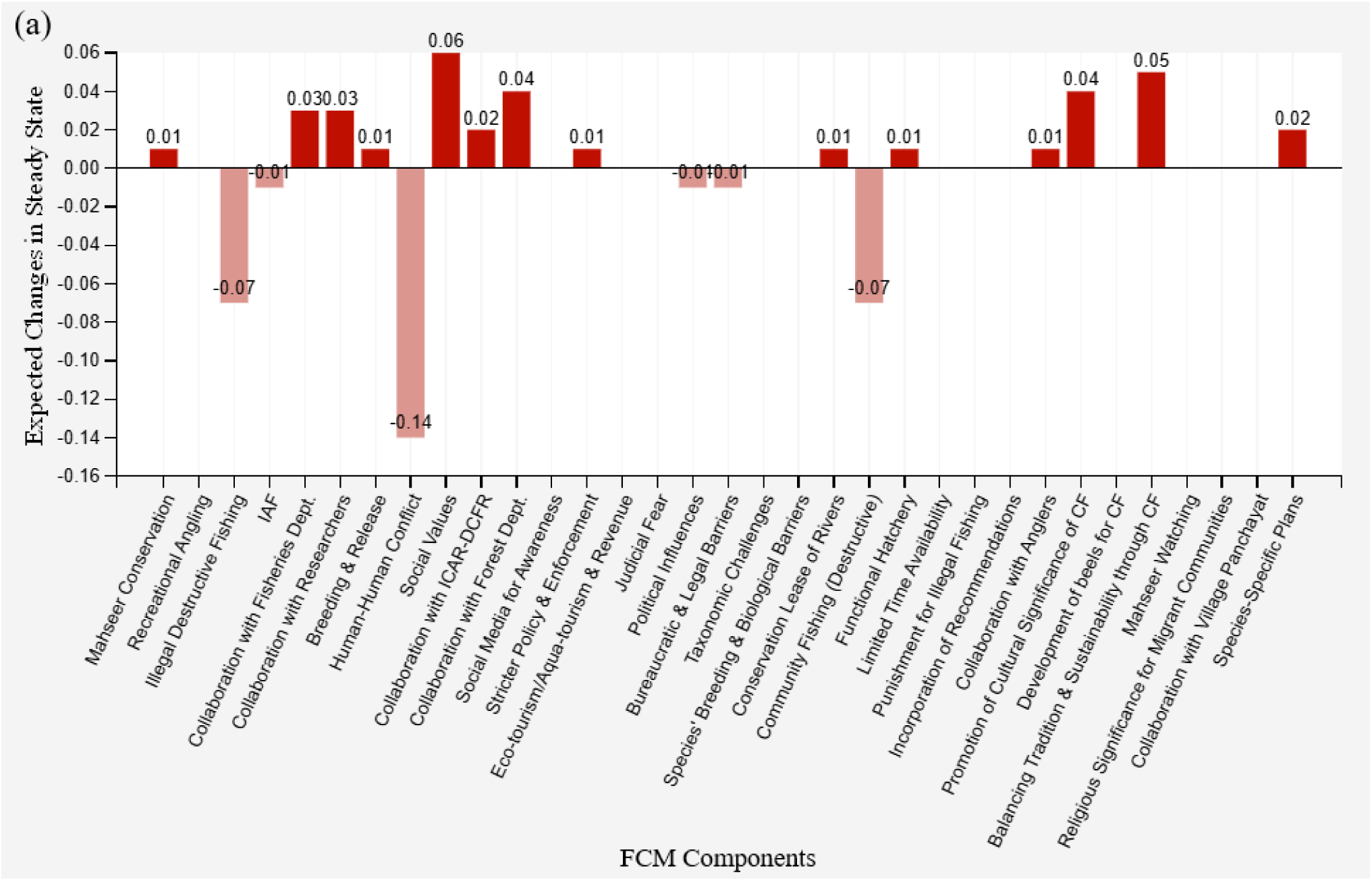
(a) Relative expected changes in the components of mahseer conservation FCM (collective) for Assam under different clamping scenarios: Scenario 1. Values above zero indicate a positive change (red bars), while values below zero indicate a negative change (pink bars). Refer to the ‘Results’ section for further details.

**Fig 8.**
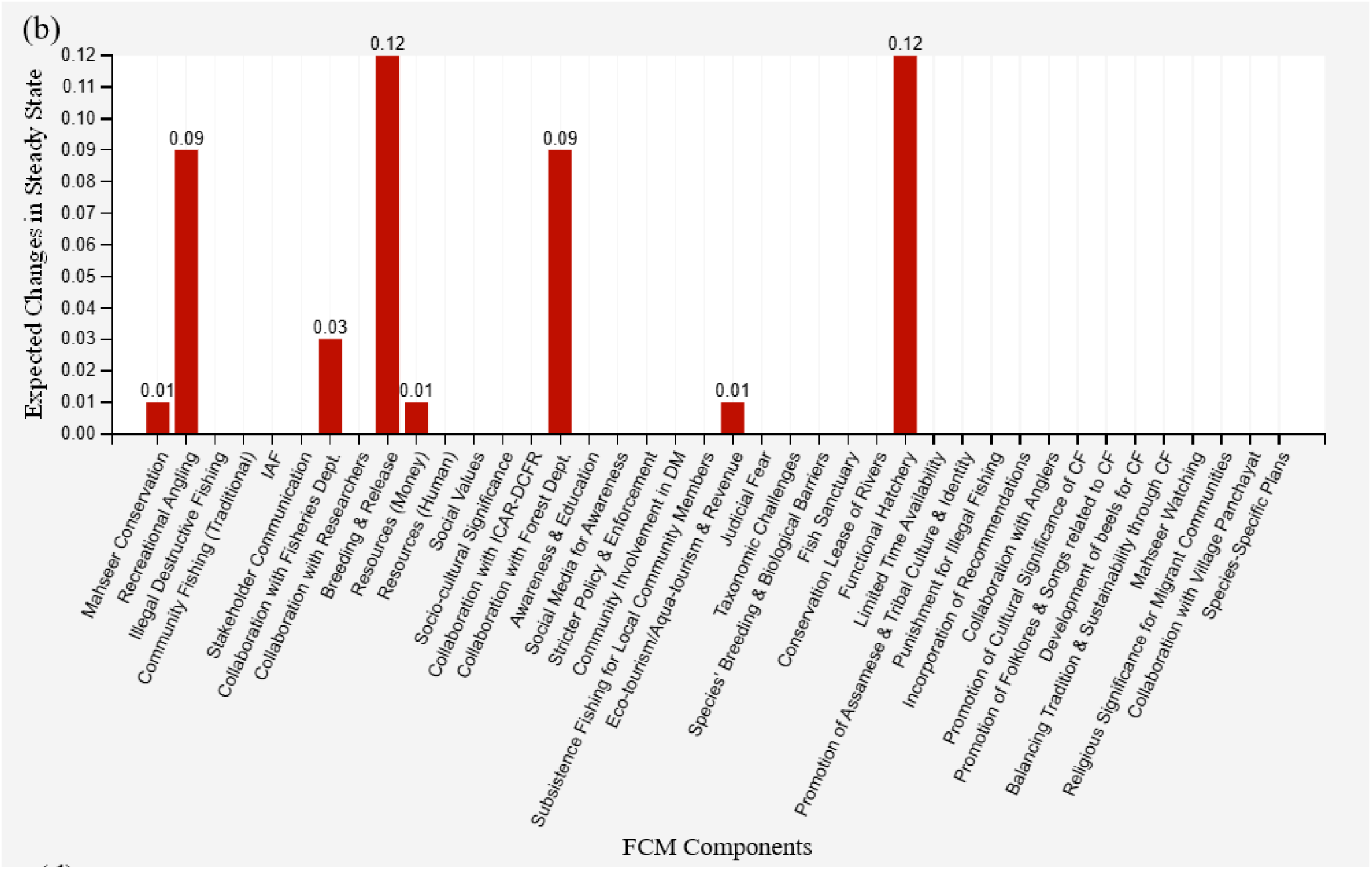
(b) Relative expected changes in the components of mahseer conservation FCM (collective) for Assam under different clamping scenarios: Scenario 2. Values above zero indicate a positive change (red bars), while values below zero indicate a negative change (pink bars). Refer to the ‘Results’ section for further details.

**Fig 8.**
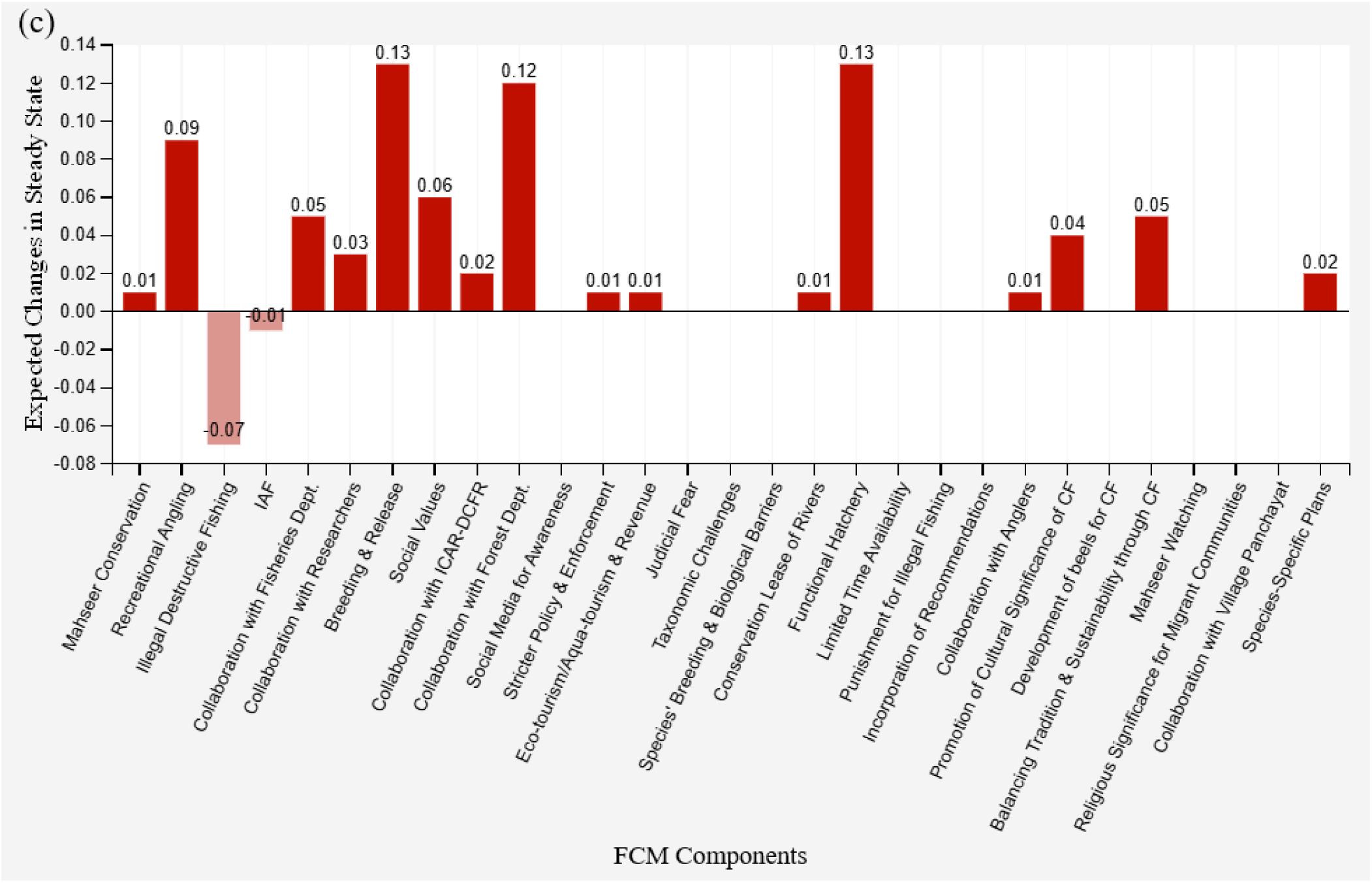
(c) Relative expected changes in the components of mahseer conservation FCM (collective) for Assam under different clamping scenarios: Scenario 3. Values above zero indicate a positive change (red bars), while values below zero indicate a negative change (pink bars). Refer to the ‘Results’ section for further details.

**Fig 8.**
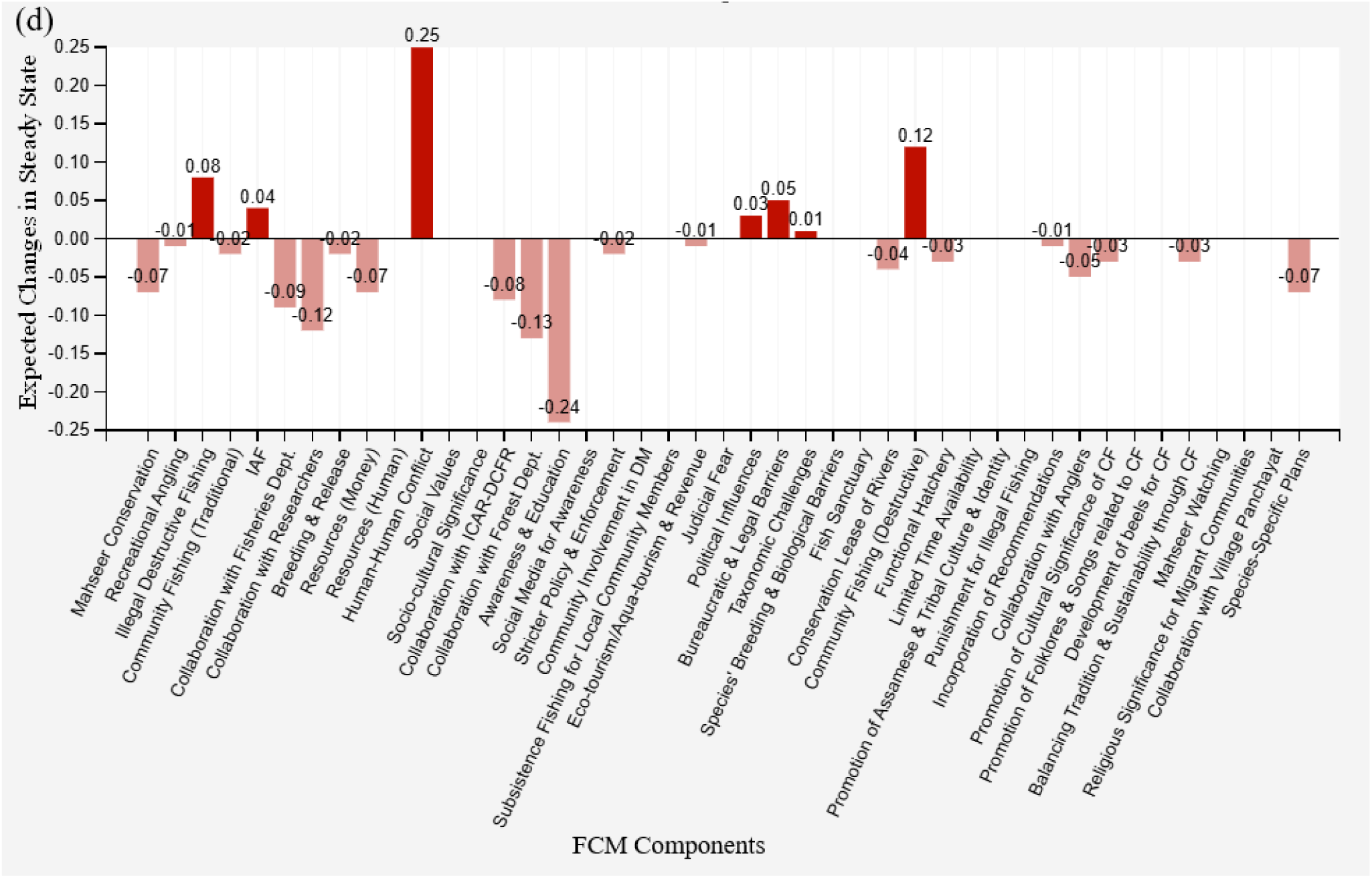
(d) Relative expected changes in the components of mahseer conservation FCM (collective) for Assam under different clamping scenarios: Scenario 4. Values above zero indicate a positive change (red bars), while values below zero indicate a negative change (pink bars). Refer to the ‘Results’ section for further details.

Scenario 2 entailed decreasing both community fishing (destructive) and human-human conflict to -1, and bureaucratic and legal barriers, and political influences to −0.5 (Fig 8b). Here, only 20% (8 out of 40) of the components showed deviation from the steady state. The breeding and release (0.12), functional hatchery (0.12), recreational angling (0.09), and collaboration with forest department (0.09) showed an upwards tendency, while collaboration with fisheries department (0.03), monetary resources and eco-tourism/aqua-tourism and revenue (all 0.01; Fig 8b) showed only a marginal increase. Furthermore, when Scenarios 1 and 2 were combined together (Scenario 3; Fig 8c), a greater proportion of the components (62.07%; 18 out of 29) moved from their steady state. Here, in addition to the changes that occurred in the components in Scenarios 1 and 2, breeding and release (0.13), functional hatchery (0.13), collaboration with forest (0.12) and fisheries department (0.05) increased further, while illegal destructive fishing (−0.07) and the presence of IAF (−0.01) decreased under this scenario (Fig 8c). However, across all three scenarios tested, ‘mahseer conservation’ showed only a marginal positive change (0.01). In Scenario 4, the most central component of the Assam collective FCM, i.e. stakeholder communication (Fig 8d) alone was decreased (-1). A shift noticed in the majority of the components (60.46%; 26 out of 43) indicates that a decline in inter-stakeholder communication would bring down multiple vital components of the mahseer conservation ecosystem such as awareness and education (−0.24), collaboration with the forest department (−0.13), researchers (−0.12), fisheries department (−0.09), and mahseer conservation (−0.07). Furthermore, in such a scenario the barriers of mahseer conservation, human-human conflict (0.25), community fishing (destructive; 0.12) and illegal destructive fishing (0.08) would acquire more strength (Fig 8d; Table 5a).

**Table 5.**
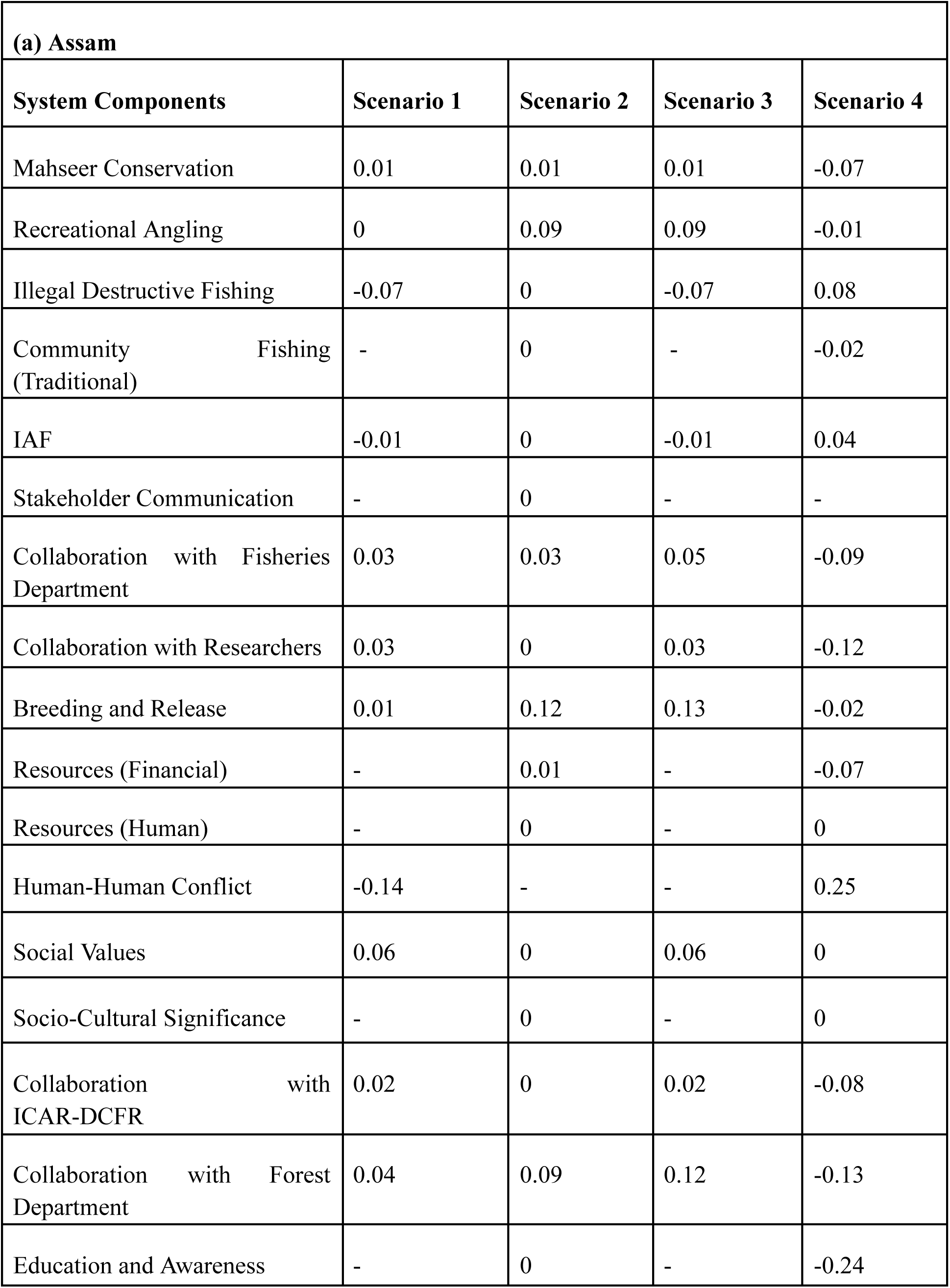
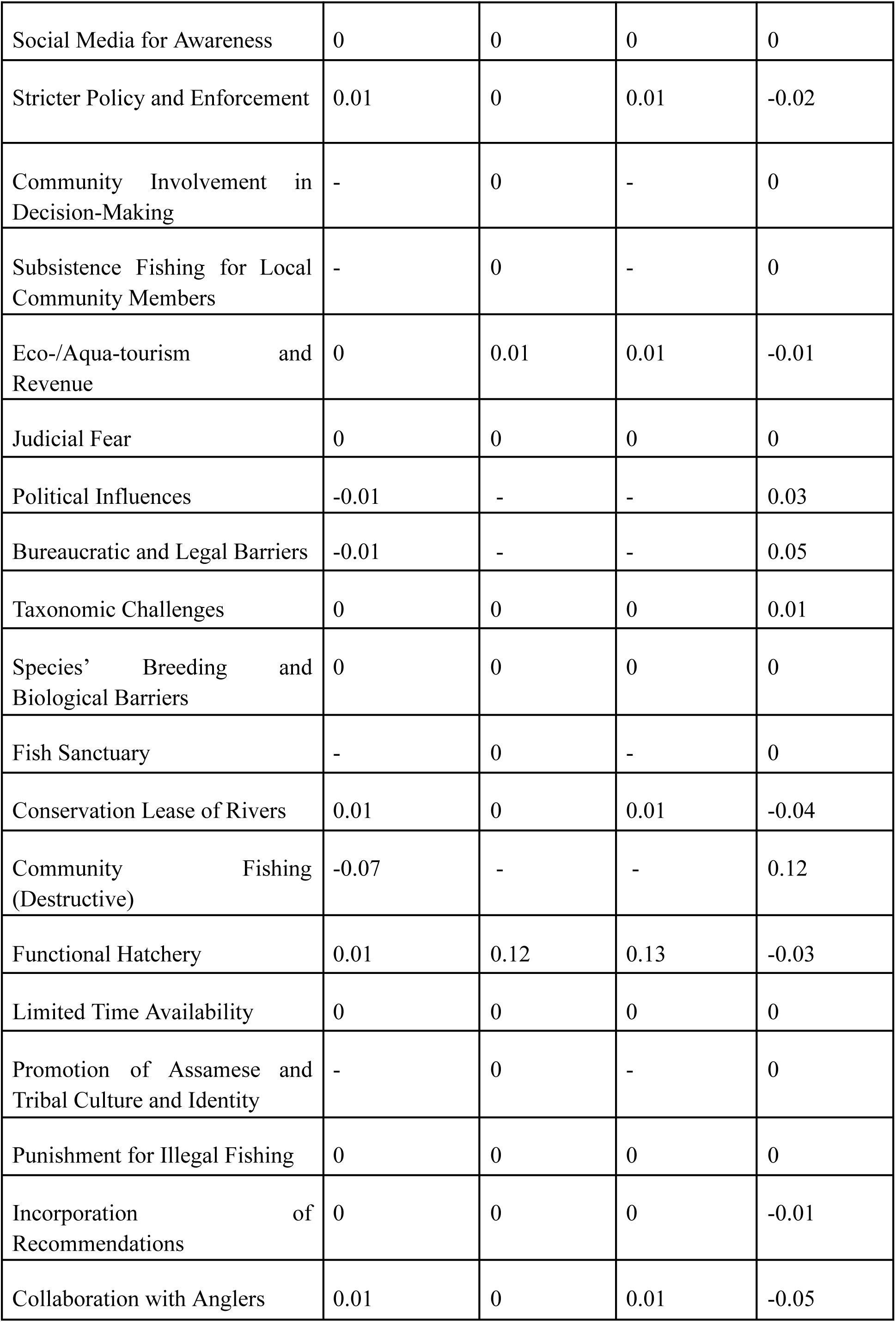
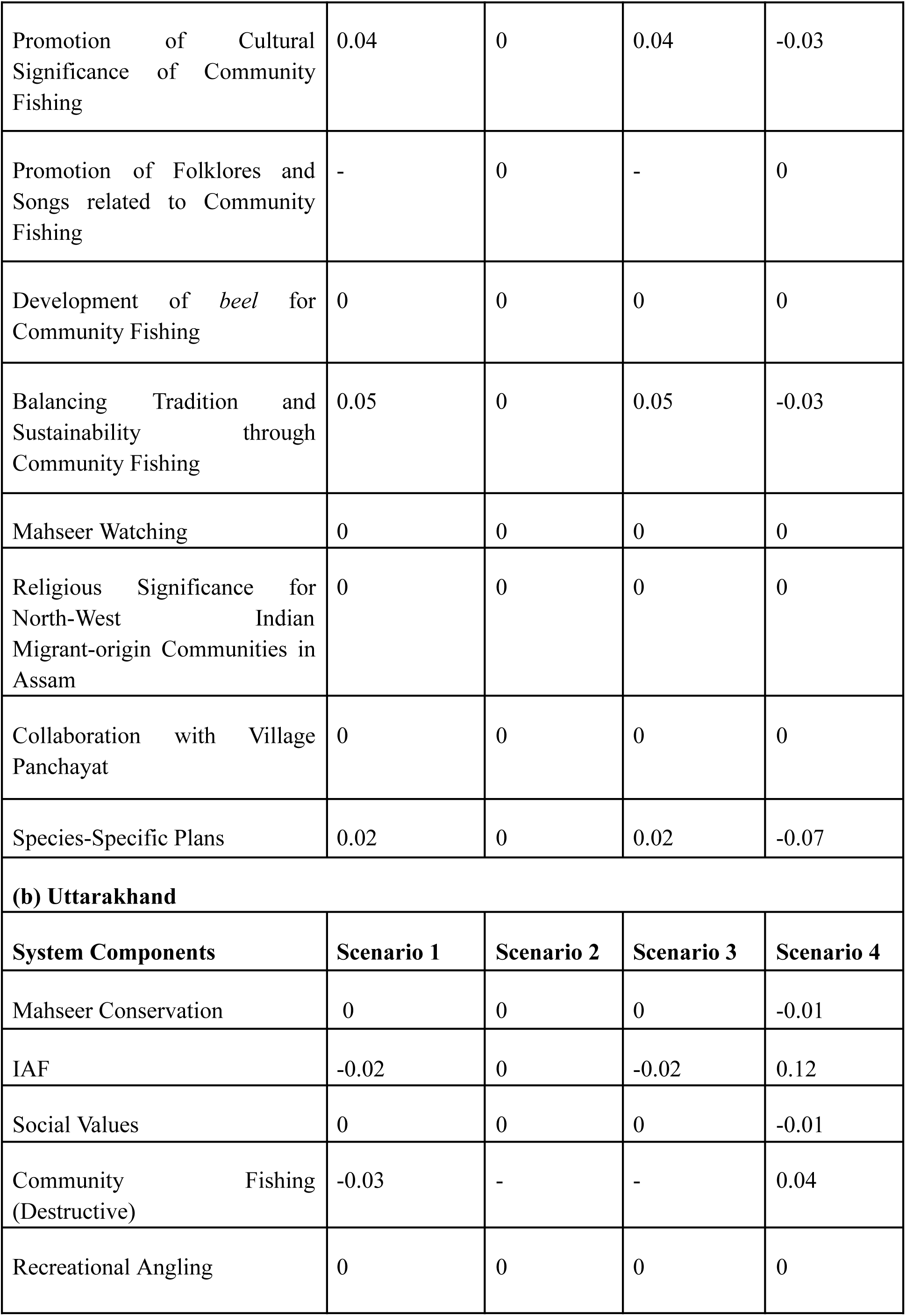
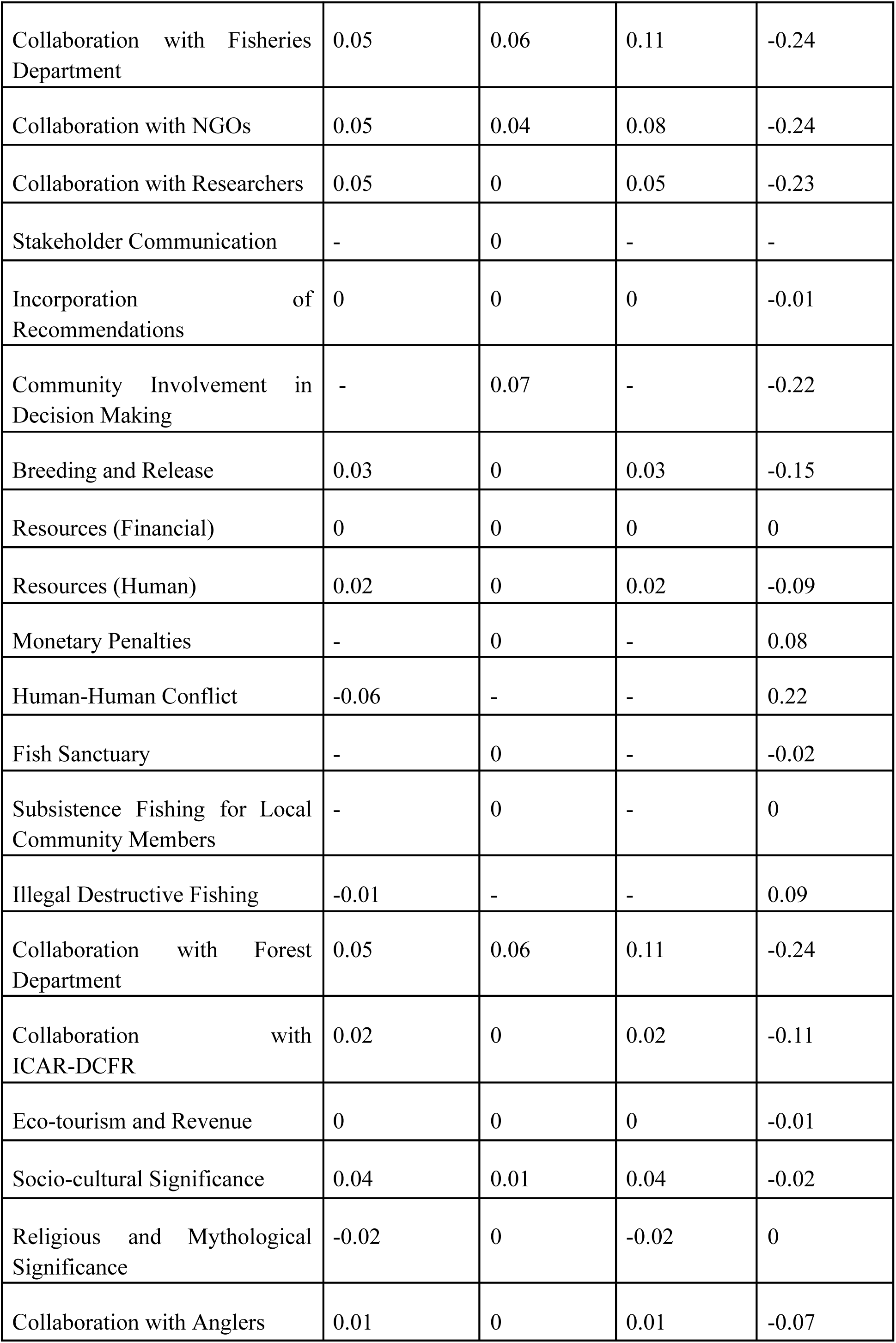
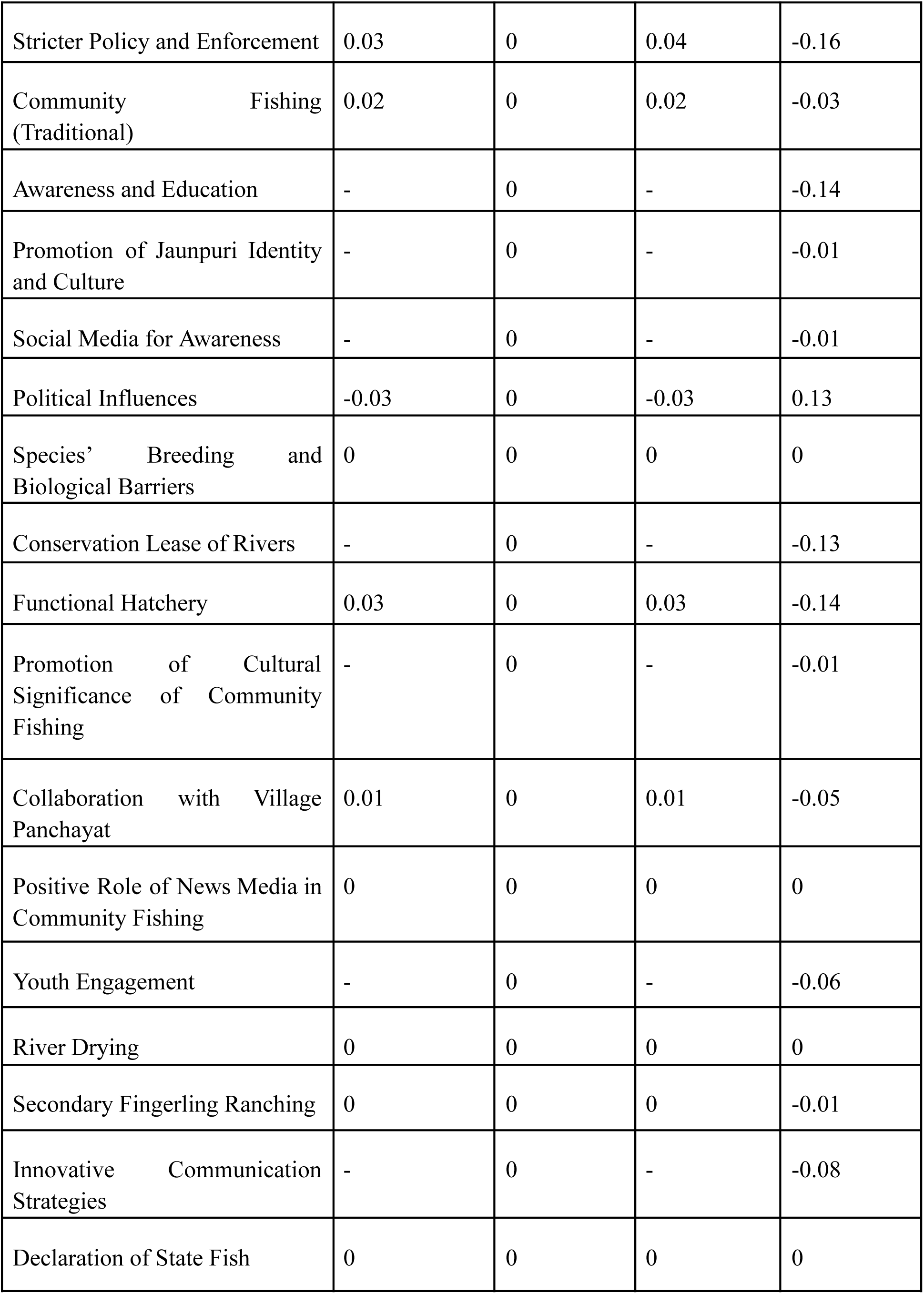
Outcomes of the ‘what if’ scenario analysis in (a) Assam and (b) Uttarakhand FCMs. Refer methodology for details.

The collective FCM of Uttarakhand was clamped as follows for Scenario 1 (Fig 9a): stakeholder communication, awareness and education, and community involvement in decision making were increased to +1. The components youth engagement, subsistence fishing for locals, promotion of Jaunpuri culture and identity, promotion of cultural significance of sustainable community fishing, monetary penalties for illegal fishing, conservation lease of rivers, declaration of fish sanctuary, use of innovative communication techniques, along with enhanced use of social media for spreading awareness, were clamped to +0.5. In this scenario, 63.34% (19 out of 30) of the FCM components displayed a noticeable shift from their steady state. Human-human conflict (−0.06), community fishing (destructive; −0.03), political influences (−0.03), IAF (−0.02), religious and mythological significance of mahseers (−0.02), illegal destructive fishing (−0.01) declined moderately. Meanwhile, the collaborations with fisheries department, NGOs, researchers, forest department (all 0.05), ICAR-DCFR (0.02), anglers and village panchayat (both 0.01) and socio-cultural significance of mahseers (0.04), breeding and release, functional hatchery, stricter policy and enforcement (all 0.03), community fishing (traditional; 0.02) and human resources (0.02) exhibited a moderate upward shift (Fig 9a).

**Fig 9.**
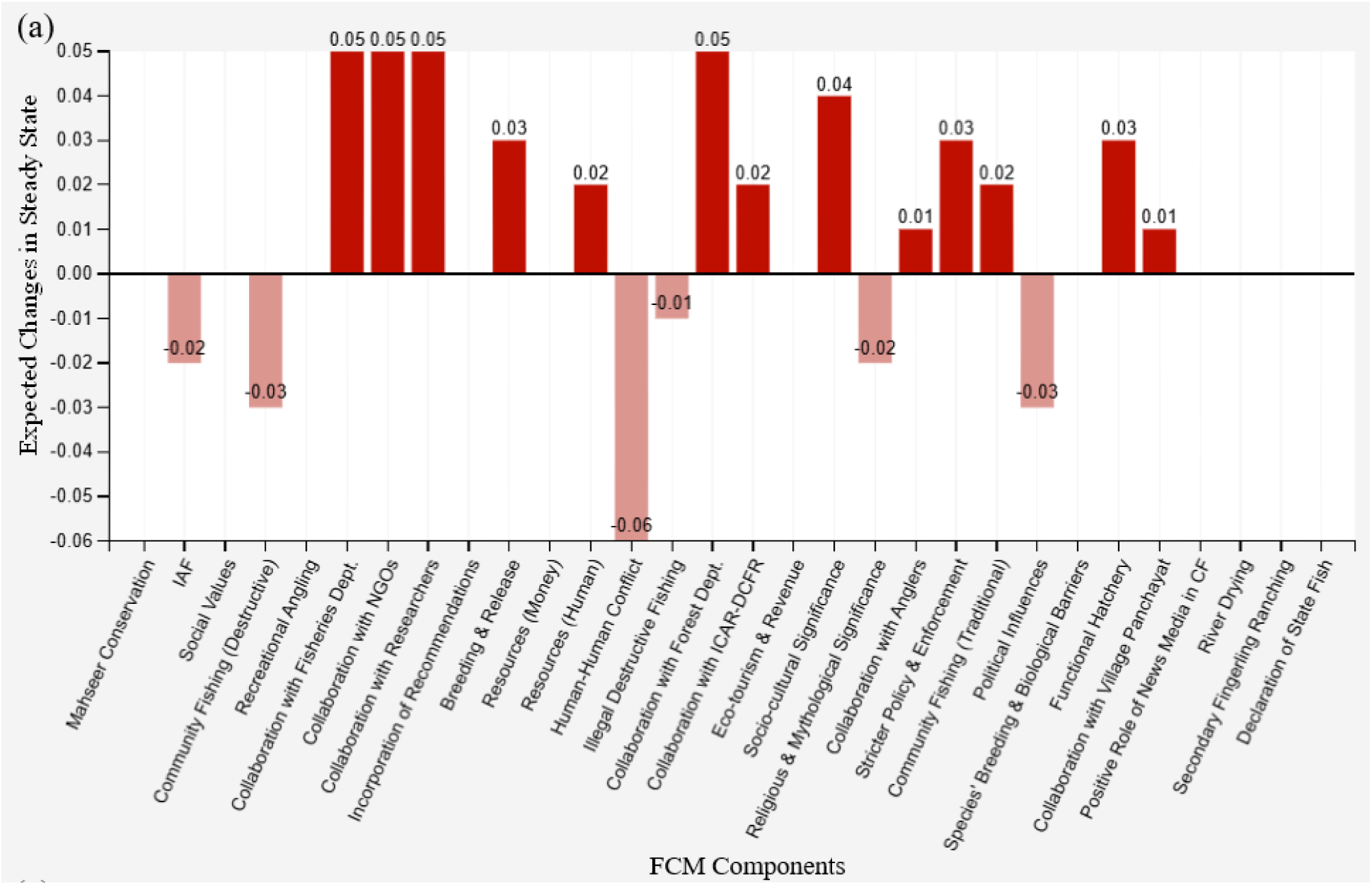
(a) Relative expected changes in the components of mahseer conservation FCM (collective) for Uttarakhand under different clamping scenarios: Scenario 2. Values above zero indicate a positive change (red bars), while values below zero indicate a negative change (pink bars). Refer to the ‘Results’ section for further details.

**Fig 9.**
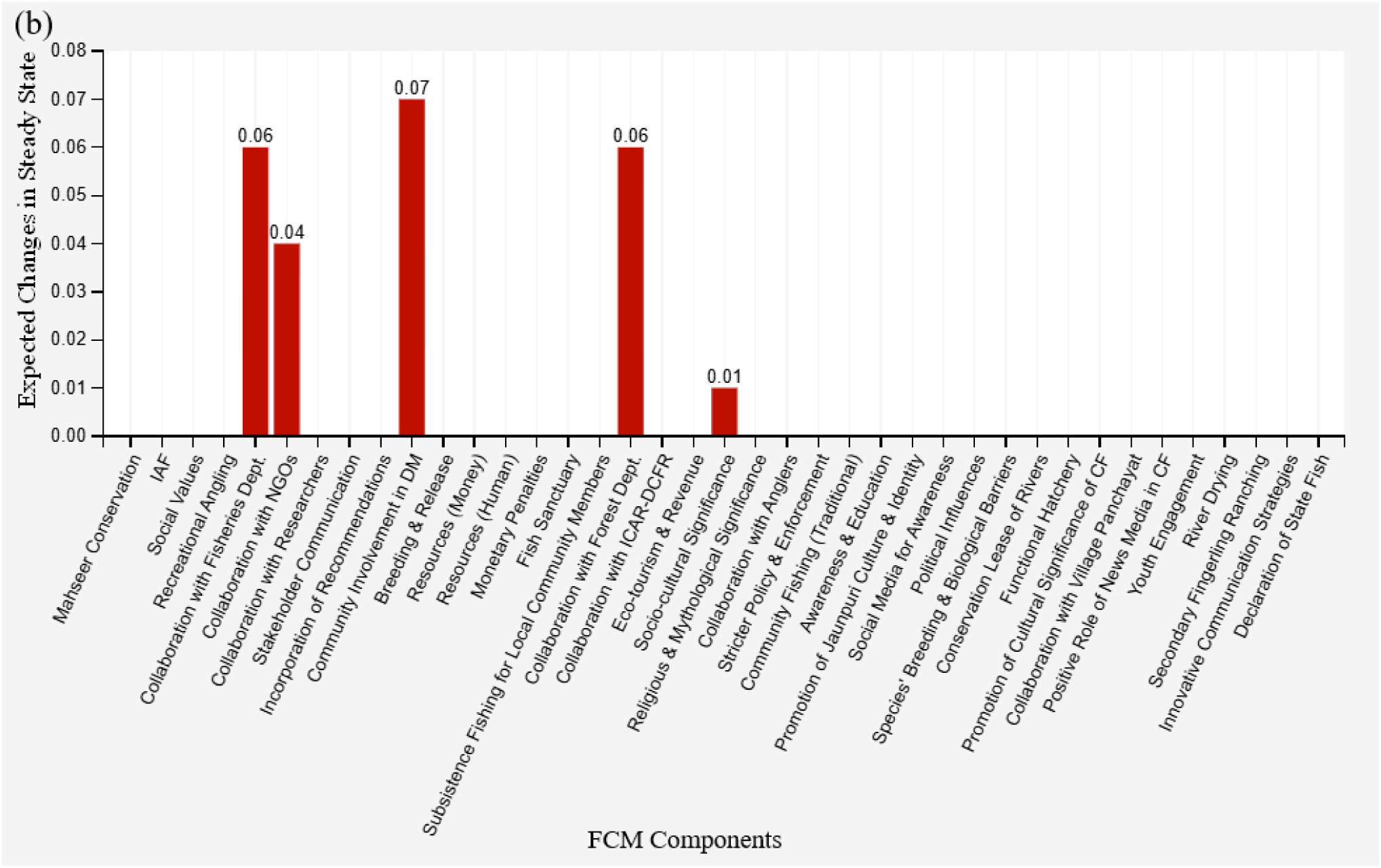
(b) Relative expected changes in the components of mahseer conservation FCM (collective) for Uttarakhand under different clamping scenarios: Scenario 2. Values above zero indicate a positive change (red bars), while values below zero indicate a negative change (pink bars). Refer to the ‘Results’ section for further details.

**Fig 9.**
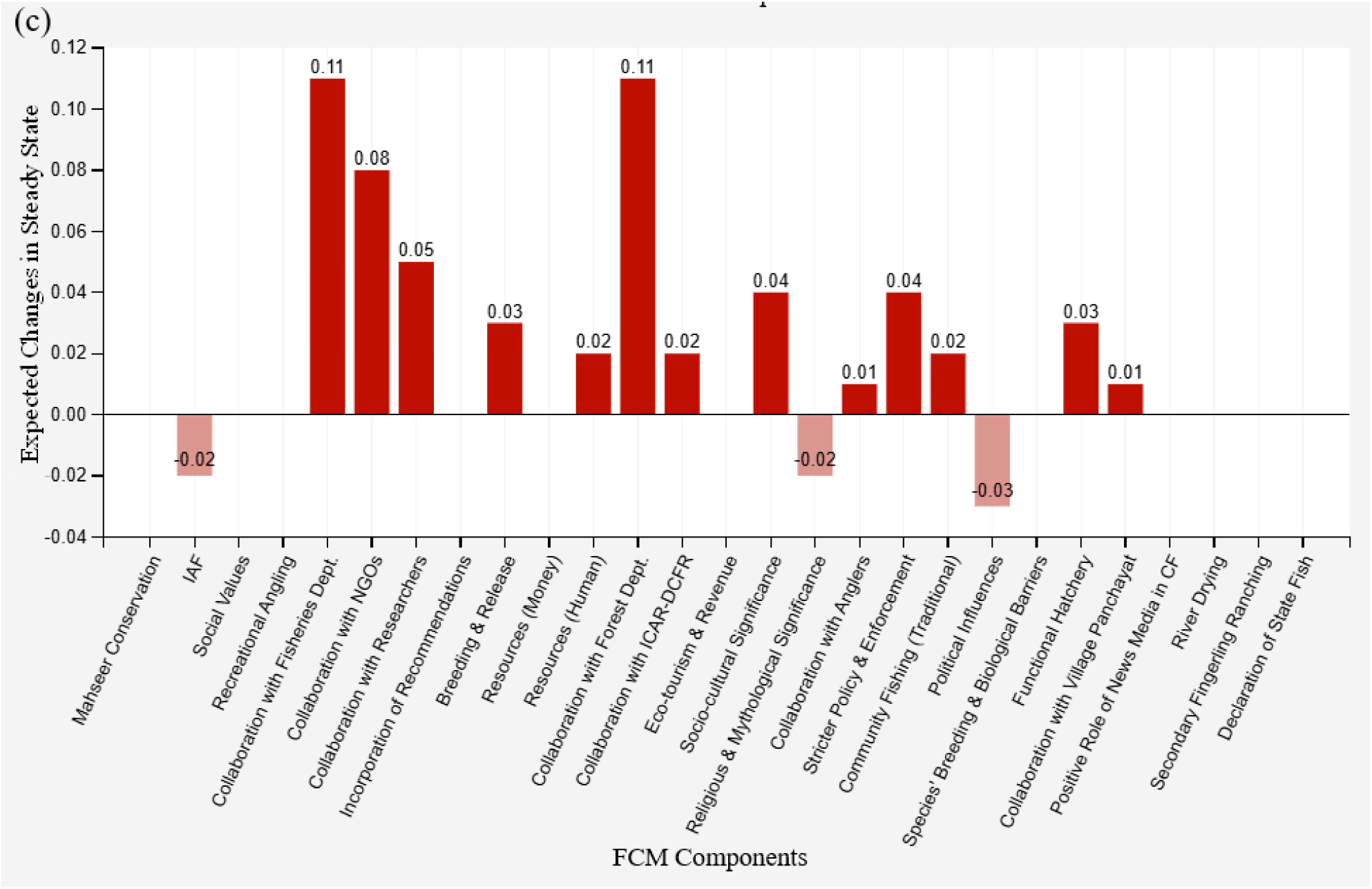
(c) Relative expected changes in the components of mahseer conservation FCM (collective) for Uttarakhand under different clamping scenarios: Scenario 3. Values above zero indicate a positive change (red bars), while values below zero indicate a negative change (pink bars). Refer to the ‘Results’ section for further details.

**Fig 9.**
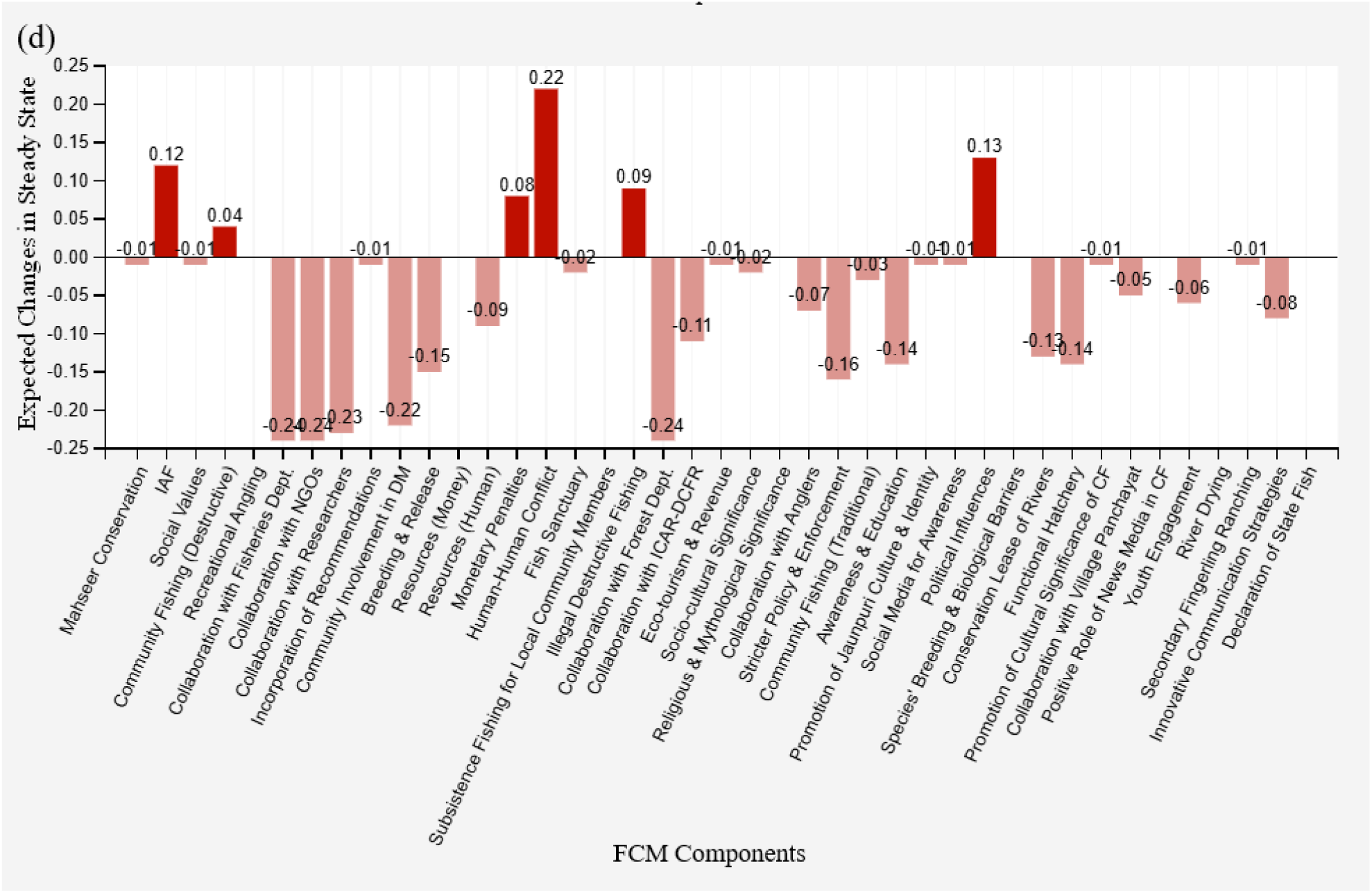
(d) Relative expected changes in the components of mahseer conservation FCM (collective) for Uttarakhand under different clamping scenarios: Scenario 4. Values above zero indicate a positive change (red bars), while values below zero indicate a negative change (pink bars). Refer to the ‘Results’ section for further details.

Scenario 2, which involved diminishing illegal destructive fishing and human-human conflict to -1, and community fishing (destructive) to −0.5 (Fig 9b) resulted in majority (87.18%; 34 out of 39) of the components remaining unchanged. The components that showed fluctuation (slightly positive) were community involvement in decision making (0.07), collaborations with fisheries (0.06) and forest departments (0.06), and NGOs (0.04), and socio-cultural significance (0.01; Fig 9b; Table 5b). Scenario 3 (combination of Scenarios 1 and 2), resulted in changes in 59.26% (16 out of 27) of the components. In addition to the components that changed under Scenarios 1 and 2, collaborations with the fisheries and forest departments (both 0.11), along with NGOs (0.08) and stricter policy and enforcement (0.04) displayed clear upward trends relative to Scenarios 1 and 2 (Fig 9c). Furthermore, three components which included political influences (−0.03), IAF (−0.02) and religious and mythological significance (−0.02) showed a mild downward trend (Fig 9c). Despite these positive changes observed in many important components, ‘mahseer conservation’ remained principally unaffected across all three scenarios tested. In Uttarakhand FCM also we tested the impact of lowering the component with highest centrality value singly, stakeholder communication, to -1 (Scenario 4; Fig 9d). A lion’s share (80.48%; 33 out of 41) of the components shifting away from their steady state in this context. Important components which exhibited decline in this scenario were collaborations with fisheries (−0.24) and forest department-0.24, NGOs (−0.24), researchers (−0.23) and ICAR-DCFR (−0.11), community involvement in decision making (−0.22), stricter policy and enforcement (−0.16), breeding and release (−0.15), education and awareness (−0.14), functional hatchery (−0.14), conservation lease (−0.13; Fig 9d). At the same time, in human-human conflict (0.22), political influences (0.13), illegal destructive fishing (0.09) and presence of IAF (0.12) increased with weakened stakeholder communication. Consequently, mahseer conservation revealed a slight downward trend (−0.01; Table 5b) in Scenario 4.

## Discussion

Many static properties of the collective FCMs, representing mental models of the stakeholders from two culturally different Indian states Assam and Uttarakhand did not show noticeable difference in many variables pointing towards a macro-level closeness in the information, attitude and actions preferred by the stakeholders (MacMillan et al. 2024; Segura et al. 2024) of mahseers habitat management and conservation. For instance, the number of components and map density of the FCM were similar for these states located in the North-Eastern and Northern regions of this geographically large nation. Furthermore, along with a higher S2 similarity index received, 32 (more than 70%) shared components and the emergence of stakeholder communication as the most central component for both states also support this argument. A lenience towards decentralised and democratic conservation mechanisms in both states is justified by the near zero hierarchy indices indicating an even and horizontal distribution of the influence across multiple components (MacDonald 1983; Gray et al. 2014). In both Assam and Uttarakhand, although no single component exhibited overwhelming control over the conservation ecosystems, stakeholder communication emerged as the central component. The highest outdegree and outdegree-to-indegree ratio received by this component in both states signify its role as the strongest driver of the mahseer conservation (Meadows 1997; Meadows and Wright 2008; Banerjee et al. 2025). Many studies are available to prove that components with higher centrality and higher outdegree-to-indegree ratios can act as the forcing functions in a system and considering their capacity to modulate other components they could be used as the leverage points for interventions (Özesmi and Özesmi 2004; Christoforou and Andreou 2017; Segura et al. 2024). Hence, improvement in the exchange of information among the stakeholders may result in better collaborations and improved work coordination resulting desired positive changes in multiple components of the mahseer conservation programmes. Engaging younger generation, particularly school and college students, in such efforts would be especially impactful as this section of the society is facing ‘extinction of experience’, but are open to novel ideas and changing their perceptions and attitude towards non-human life forms present in their environment (Binoy et al. 2021; Ian et al. 2019; Syed Azhar et al. 2022; Chellappa and Rohtagi 2025; Schmitt et al. 2025). On a positive note, many such attempts involving innovative communication strategies, youth engagement and social media campaigns were noted from many regions of Uttarakhand (Dewan et al. 2022), which needs to be replicated in other areas of mahseer conservation importance identified in the focal states. Enhancing inter-stakeholder communication can also help reduce another central component with high indegree that emerged in both states - human-human conflict (Cartwright 2006; Meierová 2020; Doley and Barman 2023). In many contexts, availability of a forum to share the ideas, plans concerns and expectation with peers and people having similar interest, discuss its implications and to reach the outcomes of such deliberation to policy makers and conservation programme managers can effectively mitigate the human-human conflicts negatively affecting the outcomes of the interventions (Nagulendran et al. 2016; Doley and Barman 2023).

In spite of these similarities, features unique to each of the focal states also emerged in the FCMs. The low Jaccard Similarity Index further supports this, indicating that although both FCMs share several similar core components, the linkages and inter-connections between them varied (van Velden et al. 2020). Moreover, although awareness and education, a variable closely related to communication, was among the top five central components in both states, in Assam, it had a high outdegree-to-indegree ratio, suggesting a more perceived deterministic role it has in the mahseer conservation ecosystem. However, in Uttarakhand, the same component resonated modulations happening in community involvement in decision making, stakeholder communication, declaration of mahseer as State Fish, youth engagement, collaborations with villagers and panchayat as well as innovative communication strategies, owing to its high indegree value. This contrast highlights the recognition of the inflow of information and its exchange by the tribal and non-tribal communities from Assam critical for achieving effective conservation outcomes by fostering desired behavioural change (Lhussier et al. 2016; Abrash Walton et al. 2022). Similarly, the component ‘community involvement in decision making’ with its high outdegree values in both FCMs announced the perceived importance of inclusivity and participatory governance (Sullivan 2019; Ullah and Kim 2021). However, in Uttarakhand the centrality of this parameter was high; any interventions empowering local communities and involving them in decision-making and governance could have broader impact in this state. The higher clustering coefficient, higher number of linkages and tightly knit component clusters, received by Uttarakhand over Assam, also support this argument since such a situation generally emerges in cognitive maps emphasising feedback loops (Gray et al. 2014). Another central component (with high in-degree) identified in Assam was the community fishing practices - both traditional and destructive - which highlight the deep socio-cultural connections local community members keep with the fishes. The strong linkages of community fishing with components such as social values, stakeholder communication, awareness and education, promotion of cultural significance, folklore and songs related to community fishing, promotion of Assamese and Tribal identity and culture indicate how these influence community fishing practices. This, in turn, reiterates the need for designing conservation programmes considering the social and cultural uniqueness of the communities cohabiting with the focal mahseer populations (Oldekop et al. 2016; Doley and Barman 2023; Jolly and Stronza 2025; Iqbal 2025).

May be stemmed from the gradual erosion of traditional systems and the growing influence of commercialisation (Singh et al. 2016; Sundriyal and Kumar 2019), ‘illegal destructive fishing’ and ‘destructive community fishing’ emerged in Uttarakhand as central components with a high in-degree value. Interestingly, in Assam FCM illegal destructive fishing did not have any prominence, so was for the traditional community fishing in Uttarakhand. In the former socio-cultural acceptance of traditional community-fishing practices, historically rooted in different tribal traditions and forms an integral component of the local identity (Chief Minister’s PR Cell, Dispur, Assam 2020; Baruah 2017; Das and Das 2020; Doloi 2024) may be helping to limiting the illegal fishing. A stronger relational values that Assamese communities attribute to mahseer and other fish populations resulting in customary norms and rituals makes regulation of such activities easier for the authorities. Although, traditional community is popular in Uttarakhand, deferring from Assam, here such events receive only limited institutional recognition, promotion and community reinforcement. The shifting socio-economic dynamics, weakening cultural cohesion, erosion of traditional socio-cultural values, and the politicisation and media promotion of non-traditional community fishing events (Sharma et al. 2016; Sundriyal and Kumar 2019; Uniyal and Uniyal 2021) add fuel to the already worsened situation in Uttarakhand. However, a few studies have also revealed many points of cultural intertwining between community fishing, popular mythologies and local identities in Uttarakhand (Kattyayani 2016; Sundriyal and Kumar 2019). Hence taking lessons and inspiration from the Assam it may be possible implement interventions through inclusive governance and enhanced communication, complemented by efforts to promote relational values, local identities, and cultural traditions (Oldekop et al. 2016; Doley and Barman 2023; Jolly and Stronza 2025) to stop the illegal fishing activities detrimental not only to mahseers but also to other indigenous aquatic life forms and their habitat.

The Assam FCM had twelve unique components; but only one from this list, promotion of folklores and songs related to community fishing emerged as a driver with significant value. Another one, bureaucratic and legal barriers, appeared also in the participant recommendations. The higher number of unique factors and drivers present in Assam, in comparison to Uttarakhand, reveals a greater number of pivot points in this state to enhance the success rate of mahseer conservation. Further analysis revealed that the five driver components with notable significance and four micro-variables with limited system-wide influence, in Assam, were mainly linked to the promotion of local culture and identity, symbols such as folklores and songs related to community fishing, involvement of local communities in decision making, and providing them the opportunities for subsistence fishing. These results point towards the need for designing decentralised and community-based conservation programs for this state, along with the biological intervention to enhance the size of the natural population (which emerged only as a micro variable with lesser effect - species’ breeding and biological barriers). Furthermore, given Assam’s rich cultural fabric and deep-rooted values (Baruah 2017; Doloi 2024), communication programmes keeping folklores and songs related to community fishing and projecting Assamese pride and identity could promote conservation awareness and sustainable fishing among locals.

In contrast, only the declaration of State Fish, which itself was a micro-variable, was identified as the driver in Uttarakhand. Here, out of the ten unique components appeared , monetary penalty for illegal fishing, youth engagement, and innovative communication techniques emerged as leverage points falling within participant recommendations. Considering these results attempts should be made to lower the judicial fear, bureaucratic and legal barriers and promote aqua-tourism centres (Das et al. 2023) through partnerships between the Fisheries Department, anglers, NGOs, and local community members. Such community-based initiatives with positive outcomes, have been reported from several regions of India. For instance, in the state Karnataka, collaborations between anglers’ associations, such as Wildlife Association of South India (WASI n.d.) and Coorg Wildlife Society (CWS n.d.), and Fisheries Department, along with local communities, have successfully fostered awareness, responsible angling practices, community participation in research, and mahseer breeding for the last many decades contributing significantly to the sustainability of natural mahseer populations. Similarly, culture-based, community-driven interventions in Meghalaya have led to the creation of multiple pristine fish sanctuaries in partnerships between government departments and local communities (Sugunan 1995; Jumani et al. 2023). Likewise region-specific factors, such as the religious and mythological significance of mahseer present in Uttarakhand (Dandekar 2013; Kattyayani 2016; Baruah et al. 2022), should be leveraged to promote the establishment of temple sanctuaries that can serve as safe habitats for the wild populations of *T. putitora*. Strengthening collaborations between news media and other stakeholders is another topic demanding immediate attention. Media attention can catalyse the efforts for making community fishing sustainable and reduce destructive fishing practices by shaping public attitudes, emotions, and perceptions (Walker et al. 2019; Trainotti et al. 2024; Das and Binoy 2025). Harnessing the wider reach social media platforms enjoys in India, along with the conventional media for positive messaging and information sharing can raise awareness and engage the younger generation, strengthen collaboration and communication among local actors (Jacobson et al. 2019; Sansui et al. 2025) and hence pursue them for actions on the ground .

The “what-if” scenario analysis in FCM offered the opportunity to simulate interventions for mahseer conservation by modulating the components that could function as the leverage points. This analysis systematically visualize ripple effects (Banerjee et al. 2025) that could happen in other components, and in many contexts in the system as a whole (Gray et al. 2013, 2015, 2019; van Velden et al. 2020; Salberg et al. 2022; Banerjee et al. 2025; Prakash et al. 2025) when selected drivers undergo decrease or increment. Subjecting driver components and recommendations with perceived positive impact on mahseer conservation (scenario 1) in Assam for this process (known as clamping) resulted in substantial reduction in destructive community fishing, illegal destructive fishing and human-human conflict, with minor gains in social values and traditions around sustainable community fishing. Nevertheless, in this scenario overall gains in mahseer conservation outcomes were limited. Scenario 2 which emphasised reducing destructive practices, bureaucratic hurdles and political interferences also produced only a marginal overall gains in mahseer conservation. However, the increase in recreational angling observed in this scenario can serve as a boon for mahseers in Assam if translated into action. Anglers in Assam, as reported from the areas around the Nameri River known for harbouring rich populations of *T. putitora,* maintain a strong ground-level connection with the local community members and have actively participated in conservation breeding of *T. putitora* in association with ICAR-DCFR (Borgohain 2015). According to Armitage et al. (2010; Raimi et al. 2022) addressing the constraints faced by government departments involved in conservation and reducing issues related to the power sharing (Jacobs et al. 2020; Riechers et al. 2025) can improve technology utilisation, inter-stakeholder collaboration and hence the on ground efficacy of conservation measures. The enhanced functional hatchery performance, breeding and release programs as well as improved collaborations with the forest department observed in the scenario resonate this point. Subsequently the third scenario representing an ideal state of increasing value of the components with positive and reducing that of the parameters with negative influence on the systems also failed to improve the ‘mahseer conservation’ noticeably.

In Uttarakhand, increasing vital components and stakeholder suggestions (Scenario 1) including communication and education, community involvement in decision-making, cultural and identity aspects, permitting subsistence fishing for locals, conservation lease of rivers and enforcing monetary penalties for illegal fishing brought about shift in more than 60% of the FCM variables. However, though any significant direct gains in mahseer conservation resulted in this scenario, a reduction in destructive community fishing practices, human-human conflicts, and political interference, and an improvement in collaboration among all stakeholders (except anglers) were observed. Clamping of this kind also supported improved breeding and release efforts, functional hatcheries, greater socio-cultural recognition of mahseers, and a revival of traditional community fishing. These outcomes align with existing evidence that social engagement and community-based local stewardship models such as the Conservation Leadership Programme (CLP) exemplified by the Baagi Village Mahseer School project in Uttarakhand (Dewan et al. 2022), can strengthen multi-actor conservation networks (Pretty and Smith 2004; Reed et al. 2014; Dewan et al. 2022). Scenario 2, which simulated reduction in illegal destructive fishing and human-human conflict, yielded no significant change in the central variable - mahseer conservation. However notable increases in community participation in decision-making as well as government departmental collaboration reveals that, reducing conflict and illegal destructive fishing activities can free up resources and conservation capacity available with the government organisations (Waylen et al. 2010; Brooks et al. 2013; Oldekop et al. 2016). Furthermore, studies on *T. putitora* in Kosi River, Uttarakhand have also underscored the importance of combining technical measures with governance interventions (Joshi et al. 2018) to sustain the mahseer populations. Akin to first and second scenario mahseer conservation outcomes did not improve, suggesting potentially deeper constraints within the Uttarakhand mahseer conservation landscape.

These results resonate with the outcome of the studies that revealed - cultural revitalisation and community involvement may mitigate harmful environmental practices, but in many contexts, it often remains insufficient to generate measurable biodiversity gains without parallel technical, ecological, organisational and/or political interventions (Pretty and Smith 2004; Beever et al. 2019). Although roles of the drivers were supported by the static analysis, results of the what if scenario points towards the need for conducting further explorations for any missing/indirect stakeholders with the potential to influence system-level leverage in both focal states. For instance, the components such as policy enforcement, funding and habitat restoration with the capability to play the central role alone and in combination with social, cultural and governance components in conservation scenarios (Cartwright 2006; Folke et al. 2010; Nagulendran et al. 2016) was not available in the Assam FCM. Similarly the role of persistent challenges such as habitat degradation and fragmentation, riverbed mining, seasonal river drying, legal ambiguities, inadequate policy implementation and enforcement (Lakra et al. 2010; Malik 2011), increasing tourism pressures, and funding gaps, and limited conservation awareness and interest among stakeholders, in deciding the future of mahseer conservation also needs to be studied in detail. Finally, scenario 4 demonstrating the cascading negative effect of diminishing inter-stakeholder communication on the systems in both states reiterated the importance of not letting the functional conservation communication programmes to weaken even a minute point while attempting for the promotion of this component. This argument is supported by a plethora of literature available revealing the role of effective communication in stakeholder - local community trust-building, promoting coordination in the activities of stakeholders, reducing human-human and conflicts destructive fishing and adaptive management of various species and ecosystems (Brooks et al. 2013; Reed et al. 2014; Beever et al. 2019). Furthermore, loss of communication networks weakening the institutional performance and conservation outcomes, and increases the chances of inter-stakeholder conflicts and frictions has also been reported (Cartwright 2006; Reed et al. 2014; Meierová 2020; Doley and Barman 2023).

The scenario analyses highlighted the complexity of mahseer conservation in Assam and Uttarakhand, where socio-cultural and ecological factors are deeply interlinked to shape outcomes (Berkes 2004; Ostrom 2009). In this context, our results emphasise the need for a holistic strategy in Assam that (i) prioritises and strengthens stakeholder communication by actively involving local community members and NGOs, (ii) integrate Assamese and tribal identity and cultural values with institutional reforms and policies, (iii) promoting a balanced blend of biological and social elements of mahseer conservation (Pretty and Smith 2004; Beever et al. 2019) and (iv) exploring additional leverage points beyond those identified in the scenarios by expanding the scope of exploration to more areas of mahseer conservation importance. In Uttarakhand (i) protecting, promoting and expanding stakeholder communication networks, (ii) strengthening youth- and community-based education and outreach through educational institutions and social media, (iii) integrating religious and mythological significance of mahseer with conservation initiatives, (iv) ensuring transparent and adaptive governance to minimise political interference and address conflicts over penalties for illegal fishing, and (v) reinforcing relevant elements of cultural identities associated community fishing can help maintaining mahseer populations. Hence, a multilayered approach integrating techno-social solutions, ecological restoration, participatory actions, adaptive and dynamic governance, evidence-based policies, embedded with mechanisms to receive and incorporate feedbacks from diverse stakeholders into the intervention programme is the need of the hour to ensure efficient mahseer conservation in both states. Once developed, such a plan could function as a model for other Indian states working to protect their wild mahseer populations.

Although our study provided many valuable insights into the stakeholder mental models of mahseer conservation in Assam and Uttarakhand, it is not free of limitations. A key inherent constraint of the FCM approach lies in its static nature; it captures stakeholder perceptions at a single point in time and cannot account for the subtleties of the participants’ perspectives, views or knowledge which is susceptible to change over time (Özesmi and Özesmi 2004). Given the dynamic nature of socio-ecological systems, the context-dependent divergences within them, and the heterogeneity in the decision-making strategies chosen by stakeholders with varying perspectives and expectations, such static representations can capture only selected dimensions of the evolving complexities of mahseer conservation. Hence, continuous evaluation and iterative updating of various components emerged in the FCM at both individual- and system-level is essential to capture the evolving trajectories of mahseer conservation. Another shortcoming of our study is the absence of information from participatory workshops conducted bringing together representatives of all stakeholder groups. Although stakeholders share, discuss and refine their mental models, as projected by some FCM studies (van Velden et al. 2020; Salberg et al. 2022; Banerjee et al. 2025) during such events giving more clarity and insights, avoiding the risk of igniting inter-stakeholder conflicts if the participants are not accommodative of the views of others is not easy (Andersson and Silver 2019). Hence we did not conduct such a workshop during the present study.

### Conservation Policy Implications

In the recent years mahseers and their conservation has gained attention in many nations falling under their range of distribution (Pinder et al. 2019; Akhtar and Ciji 2024; Baruah 2024; Iskander et al. 2024; Pongsanarm et al. 2025; Rai et al. 2025), but in India rules and legal frameworks existing for this purpose remains fragmented, and the incorporation of social and behavioural dimensions into it has not gained the momentum it demands yet. At the national level, the Indian Fisheries Act, 1897 (Act No. 4 of 1897) prohibits destructive fishing practices, regulates fishing methods and seasons, and empowers state governments to frame rules for fish protection, but this legal framework lacks species-specific provisions for the mahseer conservation. Although, many states are actively working to protect their mahseers by bringing regulations (e.g. Uttarakhand Fisheries Act, 2003 (Uttarakhand Act No. 2 of 2003); Assam Fishery Rules, 1953), lack of strict implementation of laws and continued absence of mahseer from the Schedule lists of the Wildlife (Protection) Act 1972, restricts formal legal protection and limits access to national conservation funding to save these fishes (Pinder and Raghavan 2013). In contrast, many stakeholders have over time resisted inclusion of mahseer under the Wildlife Protection Act, citing local livelihood concerns (Bhatt and Pandit 2016).

Our results from Assam and Uttarakhand offer many insights for managing lacunas existing in the policy frameworks available for mahseer conservation in India. Embedding mahseer conservation within the broader river-basin management frameworks that link water-based livelihood security with ecological restoration is essential for long-lasting, tangible outcomes in this sector. Specifically policy reforms should be designed and implemented aligning ecological measures (such as habitat restoration, regulation of riverbed mining and hydro-power projects, improved ranching methods) with socio-economic strategies such as community involvement, identity protection, livelihood diversification, eco- and aqua-tourism, and conservation-oriented angling. Linking different dimensions of mahseer conservation with different national initiatives such as the Namami Gange programme under the National Mission for Clean Ganga (NMCG; Ministry of Jal Shakti, Government of India; ICAR-CIFRI 2023) and the updated National Biodiversity Strategy and Action Plan (NBSAP; MoEFCC 2024) can improve coherence in the activities conducted by different states. For the last many years the Council of Agricultural Research (ICAR) - Central Inland Fisheries Research Institute (CIFRI), Directorate of Coldwater Fisheries Research (DCFR), and non-governmental organisations (Tata Power) have been promoting hatchery-based seed production and restocking of different mahseer species (ICAR-CIFRI 2023). Exploring reasons behind a large share of the hatchery reared juveniles not reaching the reproductive status in the natural water bodies and developing behavioural based interventions to enhance their post release survival (Varma et al. 2020) also should be given more attention by the policy makers. Considering the insufficiency of the resource available with many governmental departments for the implementation of conservation programs and law enforcement, formally recognising fishermen, NGOs, and anglers as the mahseer conservation partners and empowering them with collaborative programmes, can fill this gap and foster trust, mutual respect and responsibility among them. Establishing multi-stakeholder communication platforms bridging these stakeholders and strengthening such networks through schools, youth clubs, and local organisations, can reduce the existing awareness gaps and strengthen inter-generational stewardship and pro-conservation behaviour along with enhancing inter-departmental coordination. Such a network may also protect the quickly vanishing traditional knowledge associated with mahseers and their habitats. Moreover, promoting responsible recreational angling, community fishing events and mahseer based ecotourism under the strict surveillance of the communities and establishing more temple-linked fish sanctuaries will be not only generate livelihood for the localities depend up on mahseer but also support mahseer population management. In essence, India requires an integrated national mahseer policy that bridges ecological science, cultural heritage, and community governance, with wider scope for integrating the uniqueness of local socio-ecosystems to secure the future of mahseers and the country’s magnanimous freshwater ecosystems and the biodiversity it carries.

## Conclusion

The current study explored collective mental models - a reflection of the stakeholder perception- of mahseer conservation using fuzzy cognitive mapping in two Indian states, Assam and Uttarakhand. A striking similarity observed in the number of components, map densities, and the emergence of “stakeholder communication” as a major leverage point in FCMs of both states indicated the importance of collaboration, dialogue, and participatory governance in sustaining mahseer populations. Near-zero value of the hierarchy index observed in both FCMs suggests promotion of decentralised conservation systems catalysing collective action rather than top-down administration for this purpose in both regions. Our study also highlighted unique socio-cultural features shaping mahseer conservation priorities in each state. In Assam, community fishing, both traditional and destructive, emerged as a central component reflecting its embeddedness in local culture and identities, folklores, and traditions. Hence, intertwining ecological restoration with cultural revitalisation and community empowerment built on local pride, folk traditions and tribal identity, and permitting regulated subsistence fishing may help in protecting mahseer populations present in this state. Meanwhile, for Uttarakhand, our results propose stronger enforcement of the norms and laws, youth-led community outreach and engagement, leveraging religious and community values to foster stewardship of aquatic habitats and reducing conflicts over penalties for illegal fishing to achieve this target. The “what-if” scenario analyses reaffirmed the pivotal role of communication and collaboration in mahseer conservation in both states. Increasing stakeholder dialogues, education and awareness, and participatory decision-making triggered positive ripple effects - reduced illegal destructive fishing, diminished bureaucratic and political barriers, and lesser human-human conflicts - although direct improvements in mahseer conservation was limited. These result underscores, education, awareness and promotion of cultural identities and values alone cannot drive ecological recovery and conservation. Overall, our findings call for a techno-social conservation strategy, one that integrates ecological and population management measures with participatory action, socio-culturally informed communication, and inclusive governance to ensure sustainability of the natural populations of mahseers and their habitats.

## Author Contributions

**Prantik Das:** Conceptualisation; methodology; investigation; data curation; formal analysis; visualisation; funding acquisition; writing - original draft; writing - review and editing. **V. V. Binoy:** Conceptualisation; methodology; visualisation; writing - review and editing; supervision.

## Acknowledgments

Prantik acknowledges the Human Research Development Group (HRDG) - Council of Scientific and Industrial Research (CSIR), New Delhi, India (09/1320(0001)/2020-EMR-I) for providing the research fellowship and the annual contingency grant for field-work related expenses. He extends his sincere appreciation and gratitude to all the respondents who participated in the interviews and FGDs across districts in both Assam and Uttarakhand. He also expresses his gratitude to Mrs. Bubu Sarkar Das, Mr. Subhashis Das, Dr. Monideepa Mitra and Mr. Mahendra Singh Rawat for their assistance in establishing stakeholder contacts and data collection in Uttarakhand; and to Ms. Ankita Sharma, Dr. Achyut Malakar, Mr. Abhijit Konwar and Mr. Dhireshwar Sarma for similar support and additional assistance with Assamese translations in Assam.

## Ethical Statement

The study design was approved by the Institutional Ethics Committee (IEC) of The Trans-Disciplinary Health Sciences and Technology (TDU; Protocol Number: TDU/IEC/16E/2025/PR72) and the National Institute of Advanced Studies (NIAS; Letter Number: NIAS-EC-11/03/2022). All interviews and FGDs adhered to the required institutional ethical guidelines for conducting non-invasive human research. Interviews and FGDs were audio recorded, written notes taken and photographs captured only after acquiring oral or written informed consent from the participants. The participants were assured of the anonymity of their names, personal details and designations to protect their identity and privacy. Furthermore, additional permissions were secured wherever required and necessary. The study undertaken in the Kosi Forest Range, Ramnagar, Nainital was carried out after obtaining formal approval from the Divisional Forest Officer (DFO), Ramnagar Forest Division, Nainital, Uttarakhand (Letter Number: 2873/61).

## Data Availability Statement

In order to maintain the privacy of the respondents, the datasets used in this study are kept confidential.

Funding Human Resource Development Group (HRDG) - Council of Scientific and Industrial Research (CSIR), New Delhi (09/1320(0001)/2020-EMR-I)

## Appendix

### Appendix 1: The interview/FGD questions used for building individual FCMs

1. What are the important factors to be considered while thinking about mahseer conservation in your state?
2. The relationship existing between each of the factors you mentioned and mahseer conservation is positive or negative ?
3. In which of the following categories do you place the strength of the relationship between each of these factors and mahseer conservation - low (weak), medium (moderate) or high (strong)
4. Are the relationships among these factors - apart from their link to mahseer conservation-positive or negative?
5. In which of the following categories would you place the strength of the inter-factor relationships - low (weak), medium (moderate) or high (strong)?

## Supplementary Materials

SM 1. Definitions of mental model structural metrics used in the FCM (Özesmi and Özesmi 2004; Gray et al. 2014; van Velden et al. 2020).

| **Mental Model Structural Metric** | **Definition** |
| --- | --- |
| Components (N) | Number of variables included in the model |
| Connections (C) | Number of connections between the variables |
| Driver/Transmitter | Variables with all connections flowing out (no inward ones), hence capable of driving the system with forcing functions. |
| Receiver | Variables with only flowing in connections (influenced by other variables but influences none). |
| Ordinary | Variables with connections flowing both in and out. |
| Indegree | Cumulative strength (sum) of all edge relations entering into a variable |
| Outdegree | Cumulative strength of all edge relations originating from a variable (out-arrows). Indicates extent of variable's influence on system |
| Centrality | Sum of the indegree and outdegree values of a variable. Indicates how connected one variable is to others |
| Density | Number of connections present compared to the number all possible connections ($C/N*(N-1)$) in an FCM |
| C/N | Number of connections to number of variables in an FCM |
| Hierarchy Index | Indicates how hierarchical (top-down)/democratic (mutually interconnected) the focal system is, on a scale of 0 to 1. 1 = completely top-down, with a few components overwhelmingly controlling the system; 0 = denotes a democratic mental model of the systems with multiple connections and feedback loops. |
| Clustering Coefficient (Global) | Measures of the overall probability of nodes sharing a common neighbour are also connected to each other. Indicates the degree of overall interconnectedness or 'cliquishness' of nodes within an FCM network and also reflects how tightly integrated or well connected a system is (Bhattacharya et al. 2023). |
| S2 (Similarity Index) | Measure of the degree of similarity existing between two FCMs based on the proportion of their shared components; 0 = completely dissimilar (no shared components); 1 = identical structure (same set of components). |
| Jaccard Similarity Index (for weighted adjacency matrix of FCM) | Measure of the similarity between two FCMs based on the connections between their components. It reflects the proportion of shared weighted, directed and signed edge relationships relative to the total number of unique connections in both maps; 0 = no shared edge relationships; 1 = identical edge relationships. |
| Leverage Points | Components capable of inducing system-wide ripple effects. Includes drivers, components with high centrality and out-degree values, and participant recommendations (Salberg et al. 2022). |

SM 2: Combined adjacency matrix used for building the collective FCM (a) Assam https://docs.google.com/spreadsheets/d/1x2UoGSPdaJBpuNVp2YmCYzCt2qcQO8nHiXiUo8yWRZ8/edit?gid=0#gid=0

(b) Uttarakhand https://docs.google.com/spreadsheets/d/1x2UoGSPdaJBpuNVp2YmCYzCt2qcQO8nHiXiUo8yWRZ8/edit?gid=711711346#gid=711711346

